# Slow- and fast-contracting skeletal muscle fibers have more similar cellular and molecular contractile function at 37°C than at 25°C in older adults

**DOI:** 10.1101/2025.08.27.670366

**Authors:** Brent A. Momb, Stuart R. Chipkin, Jane A. Kent, Mark S. Miller

## Abstract

As human skeletal muscle cellular and molecular contractile properties are temperature-sensitive, the ability to perform experiments at body temperature (∼37°C) may lead to a better understanding of their *in vivo* responses and potentially their effects upon whole-muscle and whole-body performance. We quantified molecular (myosin-actin cross-bridge mechanics and kinetics) and cellular (specific tension; force divided by cross-sectional area) function in slow-contracting myosin heavy chain (MHC) I and fast-contracting MHC IIA fibers from older adults (n=13, 8 female) at 37°C and compared these to results at 25°C. MHC I fibers were more temperature-sensitive than MHC IIA fibers, showing greater increases in cross-bridge kinetics (MHC I: 4.9-8.7x; IIA: 4x) and number or stiffness of strongly-bound cross-bridges (MHC I: 86%; IIA: 34%), leading to increased specific tension in MHC I (19%), with no change in MHC IIA fibers. The expected relationship between fiber force and size (cross-sectional area, CSA) was stronger at 37°C in both fiber types, explaining 80-82% of the variance compared to 51-52% at 25°C. Specific tension was unchanged with size at 37°C in both fiber types, showing that force increases proportionally with CSA, which may be due to the increased number or stiffness of strongly-bound cross-bridges at this temperature. At 25°C, specific tension decreased with size in agreement with previous experiments. Overall, MHC I and IIA fibers at body temperature (37°C) became more analogous, including similar specific tension and closer cross-bridge kinetics, and force production was more strongly correlated with fiber size compared to a non-physiological temperature.

**Abstract Figure:** 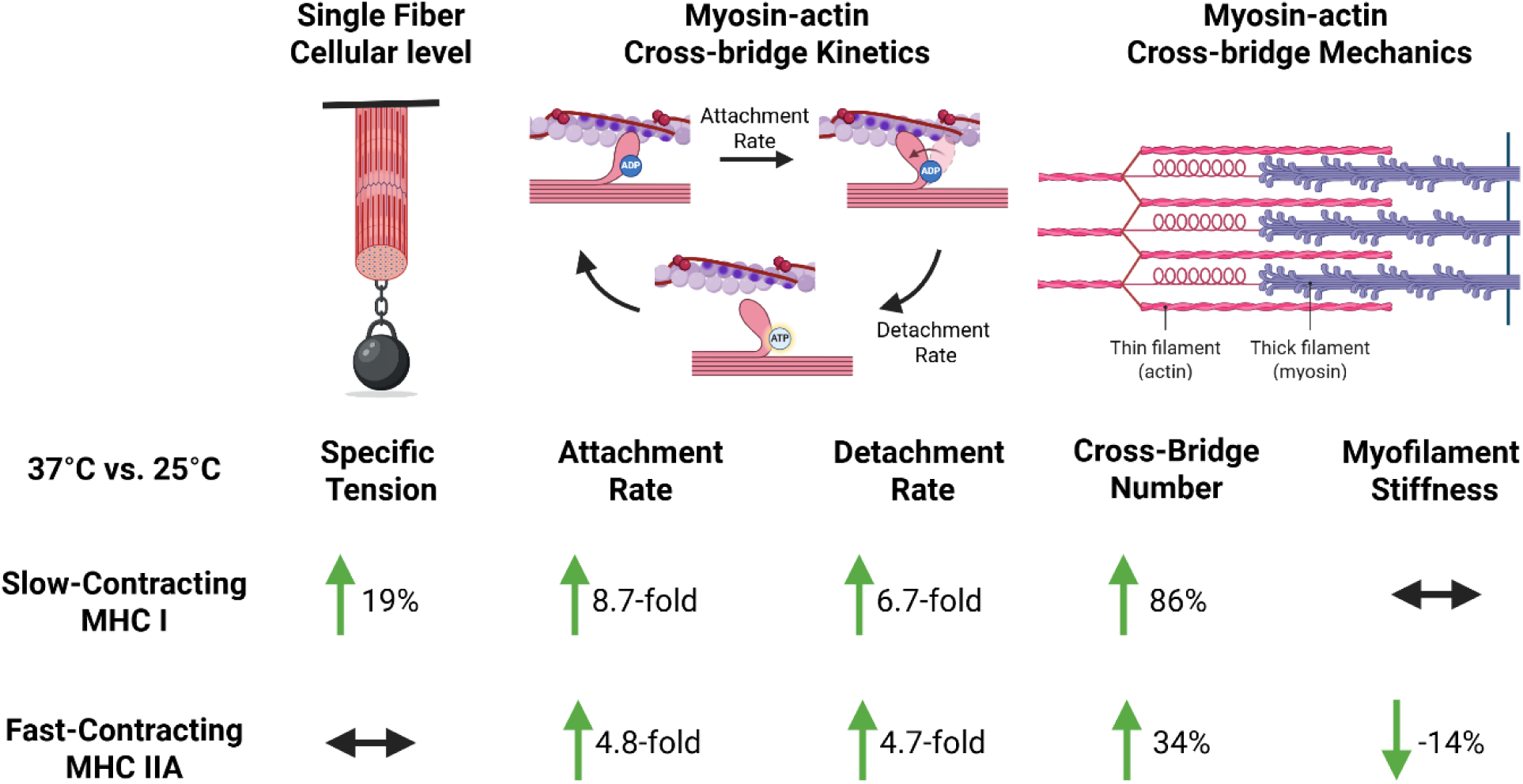

**KEY POINTS SUMMARY:**

- Although skeletal muscle function is highly sensitive to temperature, human single fiber studies have only been conducted at ≤30°C.
- Small-amplitude sinusoidal perturbations were utilized to elucidate mechanisms of single fiber force production at human physiological temperature (37°C).
- We found that functional differences in slow-contracting myosin heavy chain (MHC I) and fast-contracting MHC IIA fibers observed at 25°C were less apparent at 37°C, as force, crossbridge kinetics, and strongly-bound crossbridges increased more in MHC I fibers than MHC IIA fibers at 37 vs. 25°C.
- These results indicate that, given the different sensitivity of each fiber type to changes in temperature, functional assessments of muscle should be conducted at 37°C to better translate to *vivo* conditions.

## INTRODUCTION

Under resting conditions, human body core temperature is maintained within a narrow window of 36.3-37.7°C, with older adults presenting, on average, a slightly lower temperature by 0.23°C (Geneva *et al*., 2019). Skeletal muscle tissue temperature ranges during rest and exercise from 35-39°C (Bergh & Ekblom, 1979; Kenny *et al*., 2003; Flouris *et al*., 2015).

Performing exercise before training or competition leads to improvements in various activities, such as sprint performance, jump height, throwing distance, cycling, running, and swimming (Fradkin *et al*., 2010; Seitz & Haff, 2016), suggesting small increases in temperatures aid skeletal muscle performance. Increasing muscle temperature improves maximal rates of force development and power production(Wilson *et al*., 2025), suggesting small changes in temperature improve whole muscle contractile function. However, to our knowledge, in vitro skeletal muscle fiber function has not been examined at temperatures >30°C in humans, most likely due to increased fiber tearing or deterioration that can occur at this temperature.

Given that a primary goal of basic mechanistic studies is to infer the function of whole muscles and physical performance, not testing at actual skeletal muscle temperatures represents a significant knowledge gap, given the high temperature-sensitivity of skeletal muscle contractile velocity and power, as well as the underlying myosin-actin cross-bridge kinetics (Zhao & Kawai, 1994; Stienen *et al*., 1996; Bottinelli *et al*., 1996; He *et al*., 2000; Wang & Kawai, 2001; Debold *et al*., 2004; Knuth *et al*., 2006). When transitioning from 15°C to 30°C, slow-contracting rat fibers have a 5.6-6.0-fold increase in shortening velocity and a 19.0-26.5-fold improvement in peak power, whereas fast-contracting rat fibers display an increase of 2.3-4.0-fold for velocity and 5.1-6.8-fold in peak power (Debold *et al*., 2004; Knuth *et al*., 2006).

These studies indicate that rat single fibers have a robust increase in velocity and power with temperature that is fiber-type specific. Increasing the temperature from 25°C to 37°C increases the rate of cross-bridge attachment by 6.1x and the rate of detachment by 4.2x for slow-contracting rabbit fibers (Wang & Kawai, 2001), whereas these rates are only increased by 2.3x and 2.4x for fast-contracting rabbit fibers (Zhao & Kawai, 1994). Thus, both cellular and molecular measures in non-human models agree that contractile properties are robustly increased with temperature, but in a fiber-type specific manner, with slow-contracting fibers having a larger response than fast-contracting fibers in animals.

Human studies have shown no significant difference between the different fiber types in increases in myosin ATPase rate (Stienen *et al*., 1996; He *et al*., 2000), contractile velocity (Bottinelli *et al*., 1996; He *et al*., 2000), or power (He *et al*., 2000) with temperature (12 to 20°C). However, *in vitro* motility studies using rabbit actin filaments propelled by human skeletal muscle myosin have found that differences in actin velocity between slow-contracting (myosin heavy chain, MHC I) and fast-contracting myosin (MHC IIA and IIAX) are drastically reduced when moving from 15 to 35°C, noting that MHC IIA and IIAX velocities became similar to each other at the physiological temperature (Lionikas *et al*., 2006). Overall, these findings suggest that differences in velocity and power as well as cross-bridge kinetics between MHC isoforms may dissipate towards physiological temperatures. To adequately infer findings to whole body and whole muscle function, human single fiber contractile properties may need to be measured at *in vivo* temperatures due to their temperature-sensitivity and differential MHC isoform responses.

Not only have studies found MHC-related differences with temperature regarding contractile velocities and kinetics, but prior work has also noted the mechanisms of force production can differ between species (Kawai, 2003; Reiser *et al*., 2013). While some variation in contractile function occurs among species or type of muscle studied, isometric force is primarily due to the product of the number of strongly-bound myosin-actin cross-bridges in a half-sarcomere and the force generated per cross-bridge (Huxley, 1957; Linari *et al*., 2004; Capitanio *et al*., 2006; Palmer *et al*., 2007; Brenner *et al*., 2012). Increasing temperature has a profound impact on the underlying cross-bridge mechanics and kinetics (Zhao & Kawai, 1994), with studies noting increased force production across various MHC isoforms, ranging from slow- to fast-contracting fibers (Goldman *et al*., 1987; Ranatunga *et al*., 1987; Rall & Woledge, 1990; Zhao & Kawai, 1994; Coupland *et al*., 2001; Wang & Kawai, 2001; Coupland & Ranatunga, 2003; Debold *et al*., 2004; Knuth *et al*., 2006; Nelson *et al*., 2014; Sundberg *et al*., 2018).

Studies have been conducted to understand the underlying mechanisms behind increased specific tension (force normalized to cross-sectional area, or CSA) with temperature. These experiments have ranged from temperature jump (Goldman *et al*., 1987; Ranatunga, 2010, 2018), caged phosphate (Dantzig *et al*., 1992), length change experiments (Kuhn *et al*., 1979; Piazzesi *et al*., 2003; Decostre *et al*., 2005), and sinusoidal analysis (Zhao & Kawai, 1994; Wang & Kawai, 2001). Notably, the mechanisms for increased force generation at higher temperatures have been suggested to be either increased force per cross-bridge (Ford *et al*., 1977; Kuhn *et al*., 1979; Goldman *et al*., 1987; Dantzig *et al*., 1992; He *et al*., 2000; Piazzesi *et al*., 2003; Decostre *et al*., 2005; Ranatunga, 2010, 2018; Brenner *et al*., 2012) or increased cross-bridge number (Wang & Kawai, 2001; Kawai, 2003). Importantly, these studies have primarily been conducted on animal tissues and may not translate to humans, as others have found that human skeletal muscle may rely on both increased cross-bridge number and force per cross-bridge (He *et al*., 2000; Brenner *et al*., 2012). These prior studies call into question the ability to translate findings from lower temperatures or animal models to *in vivo* conditions in humans, highlighting the critical importance of studying human skeletal muscle tissue at 37°C.

In human skeletal muscle tissue, limitations have been encountered when attempting to assess single fiber function at 30°C, due to deterioration in MHC IIA fibers under these experimental conditions (Sundberg *et al*., 2018). These previous investigations have primarily focused on testing single fibers where substantial shortening occurs, enabling the generation of force-velocity curves. Notably, studies conducted on rabbit soleus and psoas tissue employing sinusoidal analysis have successfully extended testing conditions up to 37°C (Zhao & Kawai, 1994; Wang & Kawai, 2001). This approach uses small-amplitude, sinusoidal length perturbations under near-isometric conditions to examine cross-bridge kinetics and mechanics across various muscle types, including human cardiac (Awinda *et al*., 2020) and skeletal muscle (Miller *et al*., 2010, 2015). This technique may reduce muscle fiber damage due to shortening, as fiber length is oscillated at only 0.05% of its muscle length. Importantly, this method permits the assessment of myofilament protein function in its native three-dimensional configuration at the level of the myosin-actin cross-bridge. Under calcium-activated conditions, sinusoidal length perturbations, applied below the unitary myosin step size and across a range of frequencies (Kawai, 2003; Brenner, 2006), facilitate the determination of muscle mechanical properties relevant to specific steps in the cross-bridge cycle (Zhao & Kawai, 1993; Kawai *et al*., 1993; Mulieri *et al*., 2002; Palmer *et al*., 2007). To further improve single fiber stability, our laboratory chemically fixes the ends of the single fiber with a glutaraldehyde solution to cross-link proteins (Miller *et al*., 2010), based upon the work of others (Chase & Kushmerick, 1988; Galler & Hilber, 1998). While these techniques have been previously applied in human skeletal muscle tissue at sub-physiological temperatures of 15-25°C (Miller *et al*., 2010, 2013, 2015; Toth *et al*., 2013), their application at 37°C remains unexplored.

Considerable variability is observed when examining the specific tension of single muscle fibers, as reported in previous studies from our laboratory (Miller *et al*., 2010, 2013, 2015; Momb *et al*., 2022, 2023*b*, 2023*a*) and others (Bottinelli *et al*., 1996; Frontera *et al*., 2000; D’Antona *et al*., 2003, 2006; Gilliver *et al*., 2009; Choi & Widrick, 2010; Hvid *et al*., 2011). Because force scales with fiber size, normalization of force to CSA, or specific tension, is performed in order to compare fibers of different sizes. However, prior work has noted specific tension decreases with increasing fiber size in skinned preparations (Elzinga *et al*., 1989; Gilliver *et al*., 2009; Miller *et al*., 2015), calling into question the validity of comparing average specific tension across different populations. Some researchers have hypothesized that the diffusion of small molecules might lead to differences in the accumulation of muscle contraction by-products (ADP or phosphate), which could alter contractile mechanics (Elzinga *et al*., 1989; Stienen *et al*., 1990; Gilliver *et al*., 2009). Higher levels of by-products should affect the transition rates of the elementary steps of the cross-bridge cycle as fiber size increases, although this was not observed in our previous work in human skeletal muscle tissue at 25°C (Miller *et al*., 2015). However, because the steps in the cross-bridge cycle are highly temperature-sensitive and by-product accumulation may occur quicker at higher temperatures, further study is warranted to determine if cross-bridge kinetics are affected by fiber size.

We are not aware of any other studies that have examined the contractile mechanics of human skeletal muscle function at the physiological temperature of 37°C. The purpose of this study was to quantify single fiber contractile properties at an *in vivo* temperature and evaluate the temperature dependence of cellular (specific tension) and molecular (myosin-actin cross-bridge mechanics and kinetics) contractile functions in human skeletal muscle by comparing these measures to data collected at 25°C from the same participants. Skeletal muscle biopsies were performed on the vastus lateralis muscle of healthy older (65-80 years) participants whose habitual physical activity was measured using accelerometry.

## MATERIALS AND METHODS

### Ethical Approval

All participants provided their written, informed consent to the experimental procedures, including the skeletal muscle biopsy, which were approved by the Institutional Review Board at the University of Massachusetts-Amherst (protocol numbers 218 and 1758) and performed in accordance with the *Declaration of Helsinki*.

### Participants

Thirteen healthy older adults (65-80 years old, 8 females) completed this study. All volunteers were generally healthy by self-report, ambulatory without the use of walking aids, living independently in the community, sedentary to somewhat active (no more than two 30-minute light- to moderate-volitional exercise sessions per week) by self-report and verified by accelerometry, and females were postmenopausal (cessation of menses > 1 year). Individuals were excluded if they had a history of major neurological or neuromuscular conditions that may impact physical function; history of myocardial infarction, angina, peripheral vascular disease, surgical or percutaneous coronary artery revascularization; history of diabetes or other metabolic disease that may impact neuromuscular function; history of severe pulmonary disease, history of rheumatoid arthritis, uncontrolled hypertension (blood pressure > 140/90 mmHg), smoking in the past year, moderate to severe lower extremity arthritis, pain, muscle cramps, joint stiffness, light-headedness or other symptoms upon exertion. Individuals were also excluded based upon medications such as the use of beta-blockers, sedatives, tranquilizers, or other medication that may impair physical function, hormone replacement therapy within the preceding 5 years because this treatment may circumvent normal age-related declines in sex hormone levels, and taking anti-coagulant medication or with known coagulopathies due to increased bleeding risk from biopsy procedure. Body mass index was between 18-35 kg·m^-2^ as increased fat mass may alter single muscle fiber performance and low fast mass may be an early sign of frailty. Exclusion criteria also included any participants with a contraindication for magnetic resonance testing including a pacemaker or other implant, unintentional weight loss greater than 2.5 kg during the last 3 months, currently participating or have participated in a weight loss or exercise training program in the last year, inability to understand written and spoken English, or an inability to follow instructions, as determined by investigators during the consenting process. All participants obtained a physician’s clearance to participate. Eligibility was determined during a screening visit, where informed consent and anthropometric data were obtained.

### Accelerometry

Because habitual physical activity level can influence muscle contractile properties (Callahan *et al*., 2014; Grosicki *et al*., 2021), participants completed a physical activity log and wore an accelerometer (ActiGraph GT3X-BT) on their dominant hip for 7 days. Data were collected at 60 Hz in 1 s epochs. ActiLife software (ActiGraph, Pensacola, FL) was used to calculate moderate-to-vigorous physical activity (MVPA) using the Troiano cut points (Troiano *et al*., 2008) from a minimum of four days, including one weekend day, each of which included at least 10 hours of wear time.

### Experimental Solutions

All solutions used for biopsy processing and single fiber experiments were calculated using the equations and stability constants according to Godt and Lindley (Godt & Lindley, 1982), as described previously (Miller *et al*., 2010). Phosphate (P_i_) levels were 5 mM to correspond with resting P_i_ levels in healthy human gastrocnemius (5 mM) and quadriceps (4.5 mM) muscles (Pathare *et al*., 2005; Kemp *et al*., 2007). Characteristics of the solutions used for dissection, skinning, storage, and single fiber experiments are provided in Table 1.

**Table 1.**
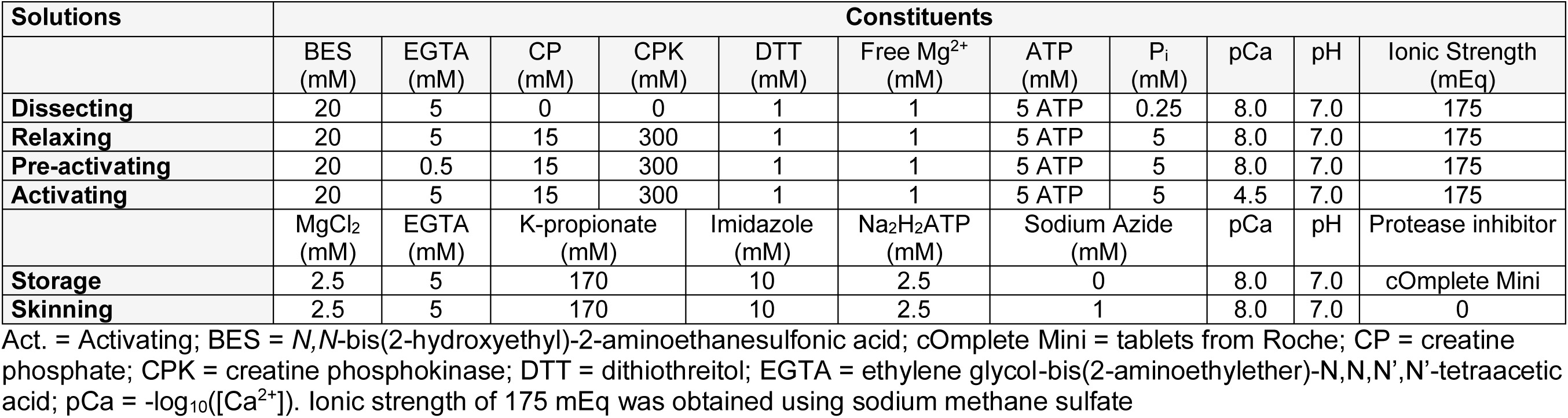
Characteristics of solutions used for skeletal muscle fibers.

### Skeletal Muscle Tissue Processing

After the accelerometry data were collected, muscle tissue from one or two legs was obtained via percutaneous biopsy of the vastus lateralis under lidocaine anesthesia with volunteers fasted overnight for at least 10 h. Most of the biopsy tissue was placed immediately into cold (4°C) dissecting solution. Muscle bundles were created by dissecting the tissue sample into bundles of approximately 50+ fibers and tying each bundle to a glass rod at a slightly stretched length at 4°C. Each tied bundle was placed in skinning solution for 24 h at 4°C and were then placed in storage solution with increasing concentration of glycerol (10% v/v glycerol for 2 h, 25% v/v glycerol for 2 h) until reaching the final storage solution (50% v/v glycerol) and incubated at 4°C for 18–20 h. Thereafter, bundles were stored at −20°C until isolation of single fibers for mechanical measurements, which occurred within 4 weeks of the biopsy.

### Preparation of Single Fibers

Single fiber preparation has been described previously (Miller *et al*., 2010). Briefly, muscle bundles were placed in dissection solution containing 1% Triton X-100 (v/v) for 30-40 min at 4°C. Segments (∼2.0-2.5 mm) of single fibers were manually isolated from muscle bundles, aluminum T-clips placed at both ends of the fiber, and the fiber was mounted onto hooks in dissecting solution at room temperature on the CSA/fixation rig. Top and side diameter measurements were made at three positions along the length of the middle 1-2 mm of the fiber using a filar eyepiece micrometer (Lasico, Los Angeles, CA, USA) and a right-angled, mirrored prism to obtain the side-to-top ratio. Fibers were then fixed at two points approximately 1-2 mm apart with glutaraldehyde, as described elsewhere (Chase & Kushmerick, 1988; Galler & Hilber, 1998), with some modifications. Fibers were placed in rigor solution (in mM: 134 potassium propionate, 10 imidazole, 7.5 EDTA and 2.5 EGTA; 5 2,3-butanedione monoxime at pH 6.8) and glutaraldehyde fixative (∼0.01% bromophenol blue (v/v), 30% glycerol (v/v), 2% glutaraldehyde (v/v)) were applied (20 s per end) using the gravity feed method (Galler & Hilber, 1998). The fiber was then placed in dissecting solution with 1% bovine serum albumin to absorb any remaining glutaraldehyde and stop it from spreading further along the fiber (Chase & Kushmerick, 1988). Any fiber remaining outside of the fixed regions was cut off along with the original T-clips. New T-clips were placed on the fixed regions, which are evident because of bromophenol blue indicator dye. The fiber was placed in dissecting solution containing 1% Triton X-100 (v/v) for 30 min at 4°C to ensure disruption of the sarcolemma and sarcoplasmic reticulum, allowing for control over the intracellular environment (e.g. contraction is initiated with high [Ca^2+^] in order to measure fiber function).

The T-clipped ends of the fiber were attached to a piezoelectric motor (Physik Instrumente, Auburn, MA, USA) and a strain gauge (SensorNor, Horten, Norway) on the sinusoidal analysis rig in relaxing solution at 15°C. The fiber was manually stretched until the sarcomere length was equal to 2.65 μm (IonOptix, Milton, MA, USA) since this resembles in vivo sarcomere length in human skeletal muscle (Chen *et al*., 2016). A camera (Point Grey, FLIR Integrated Imaging Solutions, Inc., Richmond, BC, Canada) was used to image the fiber and a computer program (ImageJ, (Schneider *et al*., 2012)) performed a fast-Fourier transformation (FFT) of the visible light and dark sarcomere patterns to determine sarcomere length. Fiber length (L) was measured using a micrometer as the distance between the inside edges of the T-clips on each end of the fiber. Fiber top width was measured in the middle of the fiber and side width was estimated using the ratio of the side-to-top width ratio obtained on the CSA/fixation rig. Fiber top and estimated side widths were used to calculate elliptical CSA.

To set the baseline tension, fibers were started in relaxing solution at 15°C, the fiber was slackened completely, the force gauge zeroed, the fiber pulled back to its original position, allowed to equilibrate for 1 min and relaxed isometric tension measured. The fiber was transferred to pre-activating solution for 30 s and then to control activating solution, with tension recorded at its plateau. After tension plateau, the fiber was returned to relaxing solution, given time to relax, and then stretched as necessary so that sarcomere length was returned to 2.65 μm. Starting again in relaxing solution, temperature was increased to 25°C or 37°C over the course of 1-2 min, and tension baseline set. At this point, single fibers were moved to pre-activating solution for 30 s and then activating solution at 25°C or 37°C, tension plateaus recorded, and small amplitude perturbation analysis performed.

### Small Amplitude Perturbation Analysis

Small amplitude perturbation analysis, or sinusoidal analysis, was adapted from previous work (Miller *et al*., 2010) and is graphically described in Figure 1 of Momb et al. (Momb *et al*., 2022). A detailed description of sinusoidal analysis (Kawai & Brandt, 1980) and how this frequency domain representation of the force response to length perturbations is related to the time domain (Palmer *et al*., 2007) have been previously published. Sinusoidal length perturbations were performed under maximal Ca^2+^-activation at 25°C or 37°C. As fiber mechanics occur at lower frequencies at lower temperatures, lower frequency sinusoidal waves must be input into muscle fibers at lower temperatures and thus experiments take longer at 25°C, ∼3 min, compared to 37°C, ∼40 s. These time frames allow for maintenance of sarcomere integrity without compromising data collection.

**Figure 1.**
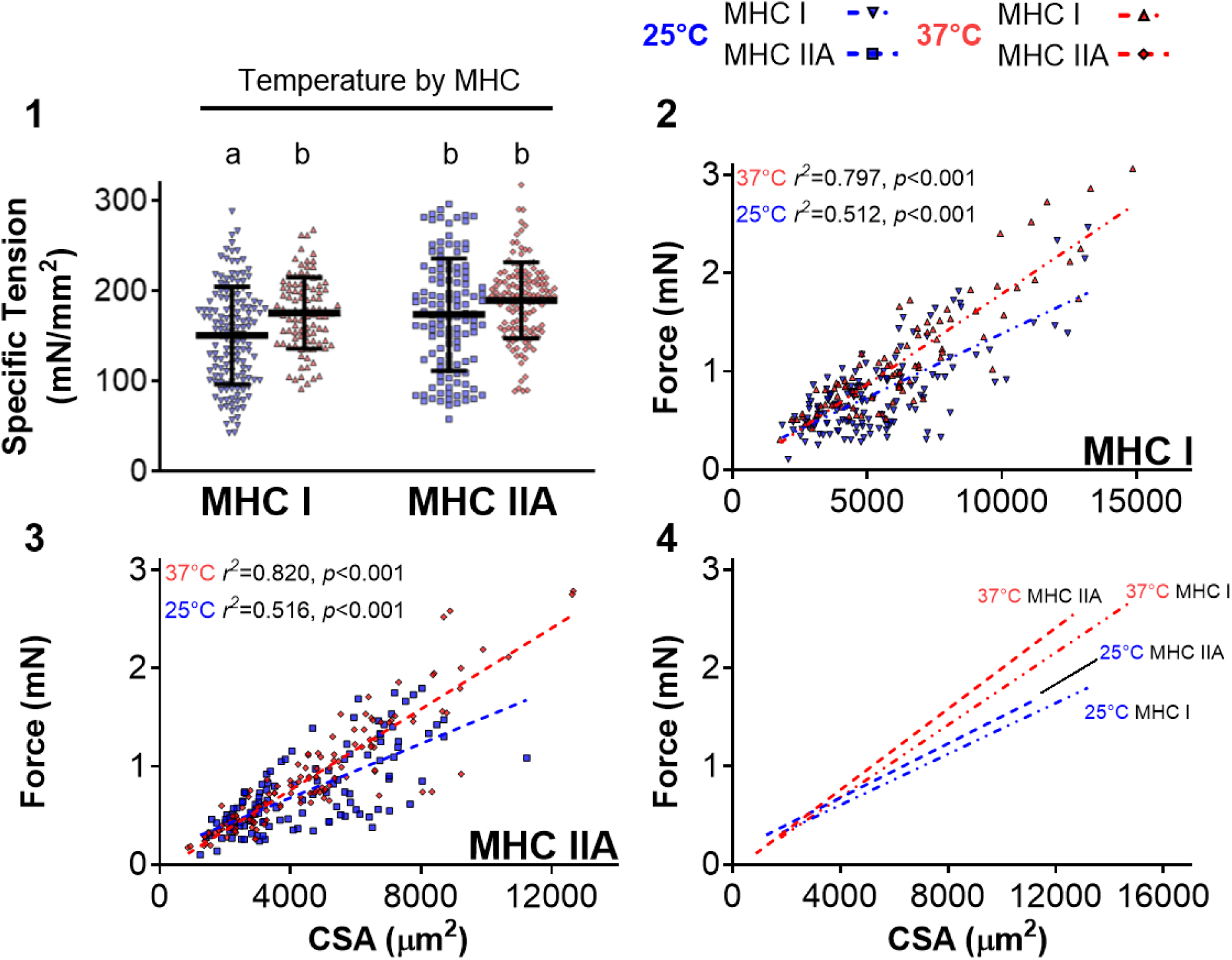
Single skeletal muscle fiber maximal Ca^2+^-activated specific tension and force versus cross-sectional area (CSA) by fiber type at 25 and 37°C. 1: Mean ± SD, with each point representing an individual fiber. Where fiber-type effects were observed, different letters above the data identify pair-wise differences (*p* ≤ 0.05) between groups. Horizontal bar indicates significant (*p* ≤ 0.1) interactions (temperature by fiber type). 2-4: Scatterplots where lines indicate linear regressions conducted for each fiber type (MHC I or IIA) and temperature. ANCOVA analysis indicated the slopes were greater and intercepts lower for both MHC I and IIA fibers at 37°C (*p* < 0.001). Slopes and intercepts for lines are as indicated: MHC I 25°C y = 1.29E^-04^x + 0.087 ; MHC I 37°C y = 1.84E^-04^x - 0.051; MHC IIA 25°C y = 0.0001.38E^-04^x + 0.128; MHC IIA 37°C y = 2.06E^-04^x - 0.065. Number of fibers tested for each group was MHC I (25°C: 147, 37°C: 98) and MHC IIA (25°C: 123, 37°C: 135).

Small amplitude sinusoidal length changes (0.05% of fiber length, or L) were applied to one end of the fiber at 48 frequencies for 25°C (0.25 to 200 Hz) or and 43 frequencies for 37°C (3 to 200 Hz). Length and force were normalized to determine fiber strain (ΔL/L) and stress (force/CSA) by dividing the length change (ΔL) by L and by dividing the force (F) by the fiber average CSA. Elastic (E_e_) and viscous (E_v_) moduli (kN/m^−2^) were calculated from the stress transient by determining the magnitudes of the in-phase and out-of-phase components (0 deg and 90 deg with respect to strain, respectively). The E_e_ and E_v_ are the real and imaginary parts (where, by definition, the imaginary part is multiplied by *i*) of the complex modulus, the ratio of the stress response to the strain. The E_e_ and E_v_ are used because these normalized values are material properties, meaning they are not dependent on CSA or L. In other words, E_e_ and E_v_ are used as skeletal muscle fibers have a variety of sizes and lengths, and normalization removes the effects of differential dimensions.

Myosin-actin cross-bridge mechanics and kinetics as well as properties of the myofilaments were derived, as previously explained (Miller *et al*., 2010) and graphically described in Figure 1 from Momb et al. (Momb *et al*., 2022), by fitting the complex modulus, meaning Nyquist plots, or E_v_ versus E_e_, were fit using the following six-parameter equation:

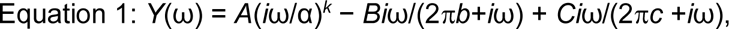

where ω=2π*f* in s^−1^, *f* is the frequency of the length perturbations (s^-1^ or Hz, frequencies used are 0.25-200 Hz for MHC I and 1-200 Hz for MHC IIA at 25°C, and 3-200 Hz for MHC I and MHC IIA fibers at 37°C), *A*, *B* and *C* are magnitudes expressed in kNm^−2^, 2π*b* and 2π*c* are characteristic rates expressed in s^−1^, i=−1^1/2^, α=1 s^−1^, and *k* is a unitless exponent. This analysis yields three characteristic processes, A, B and C, which relate to various mechanical (*A*, *B*, *C* and *k*) and kinetic (2π*b* and 2π*c*) properties of the cross-bridge cycle. The A-process (described by parameters *A* and *k*) is a linear relationship between the viscous and elastic moduli, while the B-process (described by parameters *B* and *b*) and C-process (described by parameters *C* and *c*) are semi-circles. These six parameters have been related to various aspects of muscle mechanics through experimentation and modeling (Zhao & Kawai, 1993; Kawai *et al*., 1993; Mulieri *et al*., 2002; Palmer *et al*., 2007). Although different models vary in their precise meaning, the following summarizes the laboratory’s interpretation of these parameters. The A-process has no kinetic or enzymatic dependence (Q_10_ of ∼0.9 (Mulieri *et al*., 2002)) and reflects the viscoelastic properties of the structural elements of the fiber across the oscillation frequency range. Under fully relaxed conditions, where no myosin heads are attached, the A-process represents the viscoelastic properties of the underlying fiber structure. Under Ca^2+^-activated conditions where myosin heads are attached, the A-process represents the underlying lattice structure, as well as the myosin heads attached in series (Mulieri *et al*., 2002; Palmer *et al*., 2004). The parameter *A* indicates the magnitude of a viscoelastic modulus and *k* represents the angle at which the A-process lies relative to the x-axis. Thus, *k* reflects the viscous-to-elastic modulus relationship of the A-process. The magnitude part of the B-process (*B*) is proportional to the number of myosin heads strongly-bound to actin and the cross-bridge stiffness (Kawai *et al*., 1993). The characteristic rate of the B-process (2π*b*) is interpreted as the apparent (observed) rate of myosin force production or, in other words, the rate of myosin transition between the weakly to strongly-bound state (Zhao & Kawai, 1993). For the C-process, *C* is equivalent to the number of myosin heads strongly-bound to actin multiplied by the cross-bridge stiffness (Palmer *et al*., 2007), and is therefore proportional to *B*. The characteristic frequency of the C-process (2π*c*) represents the cross-bridge detachment rate, or the inverse (2π*c*)^-1^ is the mean myosin attachment time to actin, *t*_on_ (Palmer *et al*., 2007). To examine the effects of CSA, as done previously (Miller *et al*., 2015), the magnitudes commonly normalized by dividing by CSA (*A*, *B* and *C*) were also unnormalized (*A*_CSA_, *B*_CSA_, and *C*_CSA_) to remove their direct relationship to fiber size and thus represent absolute magnitudes.

To determine the temperature coefficient (Q_10_), which describes how much a reaction rate or process changes for a 10°C increase in temperature, for 2π*b* and 2π*c*, the following equation was utilized:

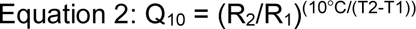

where R_1_ is the measured reaction rate at temperature 1 (T_1_ in °C) and R_2_ is the measured reaction rate at temperature 2 (T_2_ in °C). Oscillatory Work (Jm^-3^) and Power (Wm^-3^) generated by the muscle fiber were calculated from the following equations:

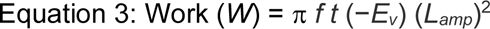

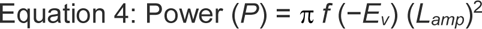

where *f* is the frequency of the length perturbations (s^-1^ or Hz), *t* is the time needed to perform the length perturbations (*s*), *E_v_* is the viscous modulus (kNm^−2^), and the fractional change in length (*L_amp_*) is 0.0005. Note that positive work and power output results from a negative viscous modulus.

### MHC Isoform Identification

After mechanical measurements were completed, single fibers were placed in a 30 ul loading buffer, heated for 2 min at 65°C and stored at -80°C until determination of MHC isoform composition by SDS-PAGE to identify fiber type as described (Miller *et al*., 2010). The stacking gel contained 4% acrylamide/bis–5% glycerol (w/v) and the resolving gel 7% acrylamide/bis– 30% glycerol (w/v). Gels ran at 70 V for 3.5 h, followed by 200 V for 20 h at 1°C. The gel was silver stained (BioRad) and the MHC isoform expressed in each single fiber was determined.

### Statistical Analysis

The temperature dependence of force and myosin-actin cross-bridge mechanics and kinetics was evaluated using a linear mixed model with group assignment as the between-subjects factor, as previously described (Callahan *et al*., 2014). A linear mixed model was used because the general linear model assumes that each measurement is independent, which is not the case for multiple fibers evaluated within each participant (i.e., fibers from the same participant are related). Accordingly, a repeated effect in the model was used to account for variations in fiber characteristics within each individual. By accounting for this within-subject variance, this approach can be considered more conservative than the process of using each fiber as an individual measurement, thus inflating sample size and statistical power. Normality was checked by evaluating the skewness, kurtosis, and the Shapiro-Wilk test. All variables assessed were normally distributed. If a main effect was noted, Fisher’s least significant difference post-hoc test was performed to determine pairwise differences between the experimental conditions. Results were considered significant at *p* ≤ 0.05. For all analyses that included potential interaction effects (temperature by fiber type), post-hoc contrasts were performed to identify pairwise differences and interactions were considered trending at 0.05 < *p* ≤ 0.10. Linear regression was conducted to examine relationships between force and other variables, such as CSA, the number of strongly bound myosin heads and/or their stiffness. ANCOVA was used to determine differences between slopes and y-intercepts for regression lines. Group data are reported as mean ± SD. All analyses were conducted using IBM SPSS Statistics for Windows version 28.0 (IBM, Armonk, NY).

## RESULTS

### Participant Characteristics

The thirteen older adults (8 female) in this study were aged 70.8 ± 1.9 yr (range 66-76), had a height of 171.4 ± 4.2 cm (range 160-178), body mass of 75.2 ± 4.5 kg (range 51.8-98.7), and body mass index of 25.6 ± 1.4 kg/m^2^ (range 18.3-31.8). Daily step count averaged 6,481 ± 2,342 (range 3,680-11,181), weekly moderate-to-vigorous activity was 282 ± 111 minutes (range 124-523), and daily activity counts were 417 ± 58 arbitrary units (range 350-539).

### Effects of Temperature on Force Production

Maximal Ca^2+^-activated specific tension (force per cross-sectional area, or CSA) was increased at physiological temperature (37°C) compared with 25°C in MHC I (*p* < 0.001), but not IIA fibers (*p* = 0.185) (Figure 1.1). The interaction effect (temperature by fiber type, *p* = 0.0534) confirms that the increase in specific tension only occurred in MHC I fibers (19%). Specific tension did not differ between fiber types at 37°C (*p* = 0.104) but was greater in MHC IIA than MHC I fibers at 25°C (*p* < 0.001). To better understand the relationship between force and CSA, force-CSA linear regressions were performed, indicating that CSA explains a greater proportion of the variance in force production at 37°C (*r*^2^ = 80-82%) than at 25°C (*r*^2^ = 51-52%) for MHC I and IIA fibers (Figure 1.2-1.4; Pearson’s correlation coefficients (*r*) in Table 2). The ANCOVA analysis indicated that 37°C increased the slopes and decreased the intercepts for both MHC I and IIA compared to 25°C (*p* < 0.001). Overall, these results indicate that force is better predicted by CSA at 37°C than 25°C.

**Table 2.** Summary of Pearson correlation coefficients (*r*) for cross-sectional area (CSA) vs. key force and kinetic variables.

| Condition | Force (mN) | Specific Tension (mN/mm<sup>2</sup>) | $2\pi b$ (s<sup>-1</sup>) | $2\pi c$ (s<sup>-1</sup>) | $A_{CSA}$ (mN) | $k$ (unitless) | $B_{CSA}$ (mN) | $C_{CSA}$ (mN) | Force/$C_{CSA}$ (Unitless) |
| --- | --- | --- | --- | --- | --- | --- | --- | --- | --- |
| MHC I, 25°C | 0.715* | -0.205^ | 0.045 | -0.084 | 0.576* | -0.194^ | 0.597* | 0.650* | -0.014 |
| MHC IIA, 25°C | 0.718* | -0.230^ | -0.077 | 0.071 | 0.594* | -0.161 | 0.636* | 0.683* | -0.036 |
| MHC I, 37°C | 0.893* | -0.031 | 0.000 | 0.001 | 0.557* | -0.044 | 0.768* | 0.734* | -0.029 |
| MHC IIA, 37°C | 0.905* | 0.104 | 0.045 | 0.039 | 0.691* | -0.008 | 0.792* | 0.774* | 0.077 |

CSA, fiber cross-sectional area; MHC, myosin heavy chain. ^ indicates significance ( $p < 0.05$ ); \* indicates significance ( $p < 0.001$ )

Maximal Ca^2+^-activated isometric force for MHC I fibers was 1.04 ± 0.56 mN at 37°C and 0.81 ± 0.41 mN at 25°C *(p* < 0.001). In MHC IIA fibers, isometric force was 0.89 ± 0.55 mN at 37°C and 0.77 ± 0.40 mN at 25°C (*p* = 0.0211). MHC I fiber CSA was 5905 ± 2138 μm^2^ at 37°C and 5500 ± 1964 μm^2^ at 25°C (*p* = 0.136). CSA of MHC IIA fibers was 4607 ± 1947 μm^2^ at 37°C and 4395 ± 2229 μm^2^ at 25°C (*p* = 0.419).

### Effects of Temperature on Elastic and Viscous Moduli

Nyquist plots of the combined elastic and viscous moduli show the effects of temperature on these curves for MHC I (Figure 2.1) and IIA (Figure 2.2) fibers. To better understand the frequency response of these parameters, the elastic and viscous moduli are individually plotted versus oscillatory frequency (Figures 2.3 and 2.4), which show that the higher temperature shifts the response of these parameters to higher frequencies. For instance, 37°C versus 25°C had a higher frequency of the minimal values of the elastic (MHC I: 20 ± 3 vs. 2 ± 1 Hz, *p* < 0.001; MHC IIA: 35 ± 3 vs. 7 ± 2 Hz, *p* < 0.001) and viscous (MHC I: 10 ± 2 vs. 1 ± 0 Hz, *p* < 0.001; MHC IIA: 15 ± 3 vs. 4 ± 1 Hz, *p* < 0.001) moduli, as well as maximum viscous modulus values (MHC I: 55 ± 5 vs. 7 ± 2 Hz, *p* < 0.001; MHC IIA: 95 ± 9 vs. 20 ± 3 Hz, *p* < 0.001) indicating faster myosin-actin cross-bridge kinetics. The oscillatory work and power as well as the frequencies at which these values occurred also increased with temperature in MHC I and IIA fibers (Figures 2.5 and 2.6, Table 3). Notably, at 37°C, the work and power output curves of MHC I and IIA fibers overlap substantially, while at 25°C these two curves are separate for these fiber types, indicating that work and power output of slow- and fast-contracting fibers are more similar at physiological temperature.

**Figure 2.**
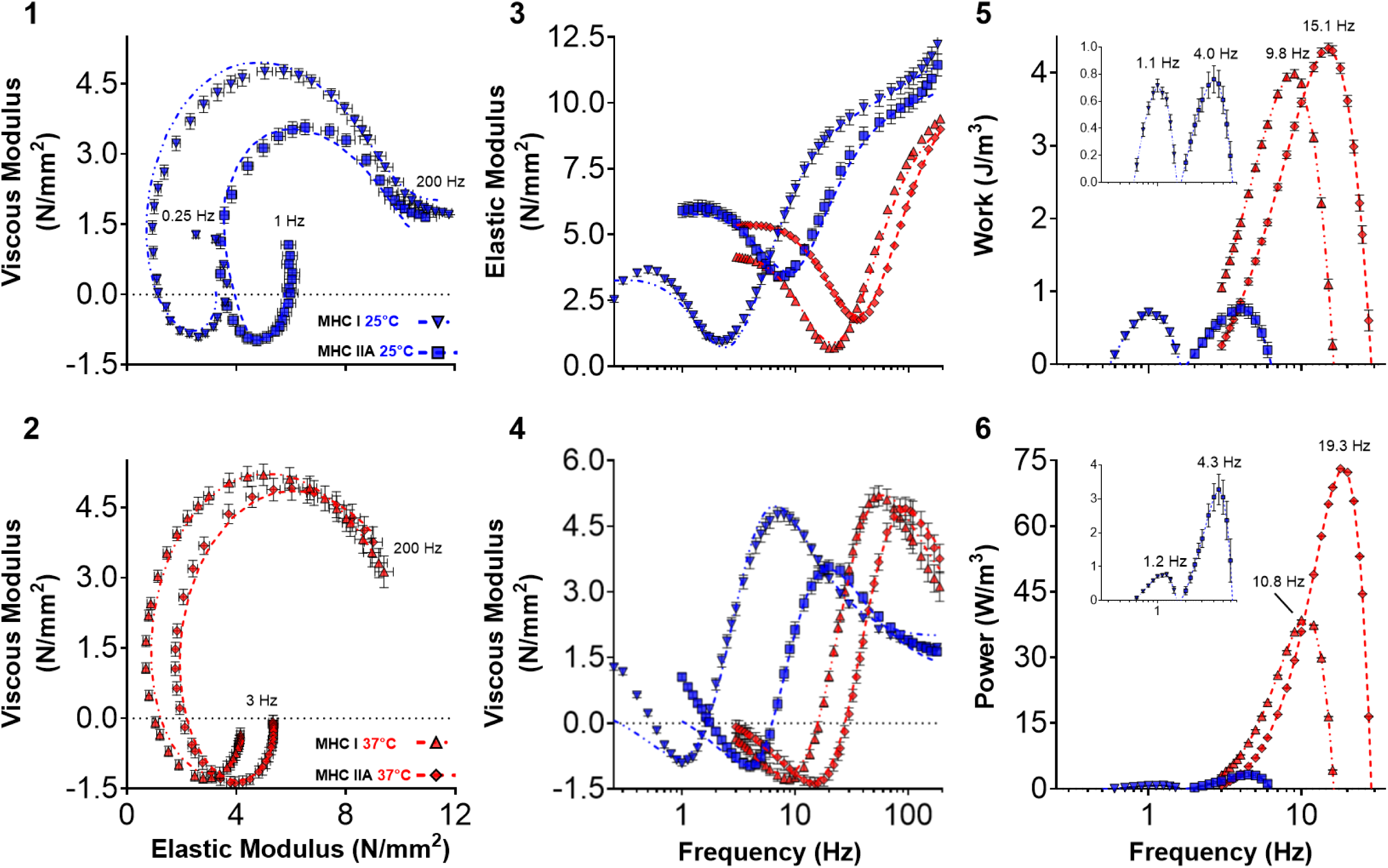
Single skeletal muscle fiber maximal Ca^2+^-activated sinusoidal analysis Nyquist plots, Bode plots, and oscillatory work and power across frequency by fiber type for 25 and 37°C. Mean ± SD for each frequency are shown. (1-4): Lines represent the fits from the 6-parameter model. Total number of fibers tested for each condition was MHC I (25°C: 147, 37°C: 98) and MHC IIA (25°C: 123, 37°C: 135).

**Table 3.** Summary of oscillatory work and power.

| Condition | n | Peak Work (J/m<sup>3</sup>) | Frequency of Peak Work (Hz) | Peak Power (W/m<sup>3</sup>) | Frequency of Peak Power (Hz) |
| --- | --- | --- | --- | --- | --- |
| **MHC I, 25°C** | 147 | 0.7 ± 0.1<sup>#</sup> | 1.1 ± 0.1<sup>*</sup> | 0.8 ± 0.1<sup>*</sup> | 1.2 ± 0.5<sup>*</sup> |
| **MHC IIA, 25°C** | 123 | 0.8 ± 0.2<sup>#</sup> | 4.0 ± 0.2<sup>*</sup> | 3.3 ± 0.5<sup>*</sup> | 4.3 ± 0.7<sup>*</sup> |
| **MHC I, 37°C** | 98 | 4.0 ± 0.1<sup>#</sup> | 9.8 ± 0.3<sup>*</sup> | 38.6 ± 0.7<sup>*</sup> | 10.8 ± 0.3<sup>*</sup> |
| **MHC IIA, 37°C** | 134 | 4.3 ± 0.1<sup>#</sup> | 15.1 ± 0.7<sup>*</sup> | 73.1 ± 0.1<sup>*</sup> | 19.3 ± 0.8<sup>*</sup> |

Data are mean ± SD; MHC, myosin heavy chain. \* indicates significant difference ( $p < 0.001$ ) from other three conditions, # indicates significant difference ( $p < 0.05$ ) between temperatures

### Effects of Temperature on Molecular Function

MHC IIA fibers had faster cross-bridge kinetics than MHC I, regardless of temperature (both *p* < 0.001). Physiological temperature increased cross-bridge kinetic rates 4.0-8.7-fold. Specifically, the rate of myosin transition between the weakly- and strongly-bound states (2π*b*) and the rate of myosin detachment (2π*c*) increased in MHC I and IIA fibers at 37°C compared with 25°C (*p* < 0.001) (Figure 3). Both 2π*b* and 2π*c* had interaction effects (temperature by fiber type, both *p* < 0.001), as these rates increased more in MHC I than MHC IIA fibers (2π*b*: 9.4-fold vs. 4.8-fold *p* < 0.001; 2π*c*: 6.7-fold vs. 4.7-fold*, p* < 0.001). These data show that cross-bridge kinetics between fiber types became more similar at physiological temperature.

**Figure 3.**
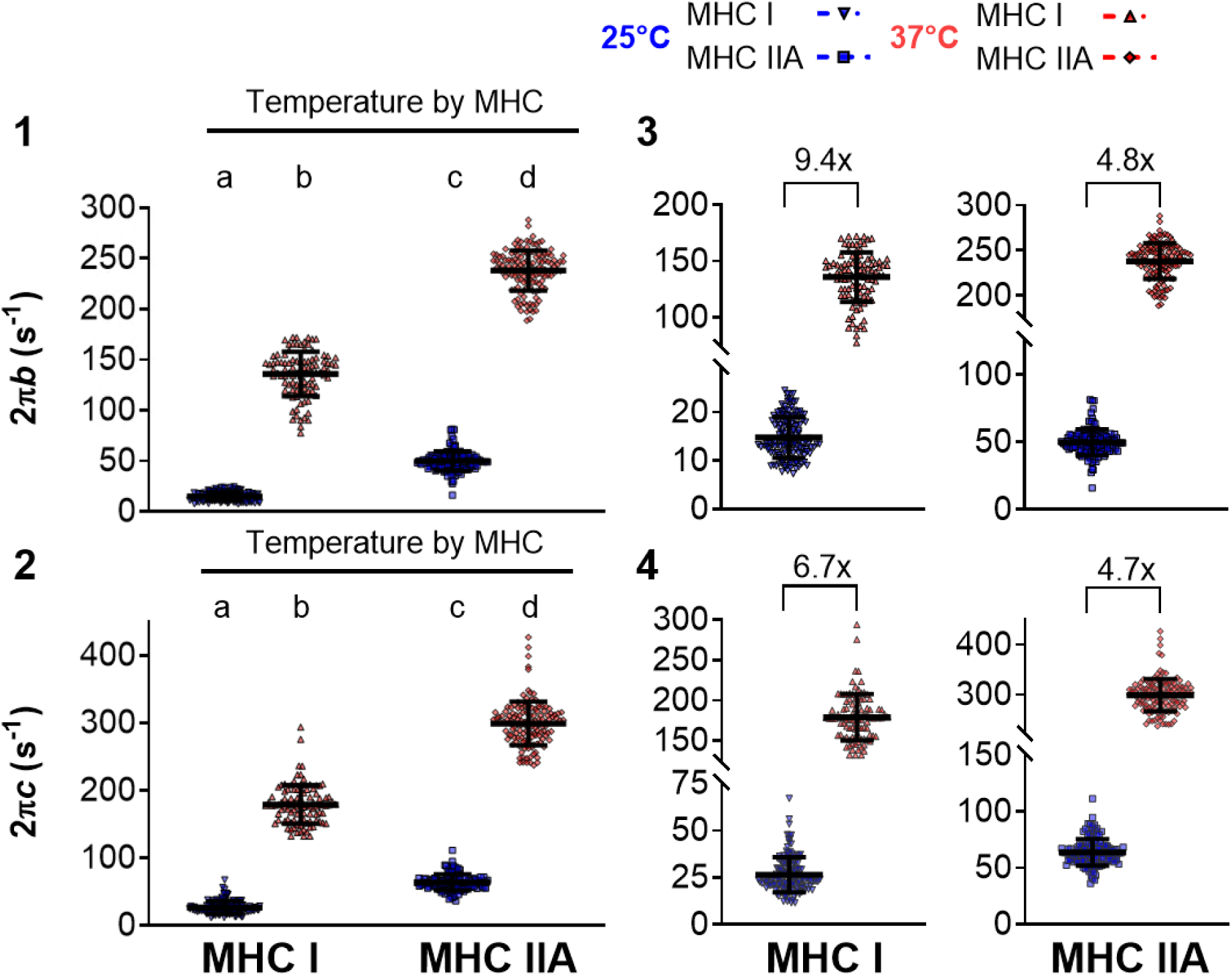
Single skeletal muscle fiber maximal Ca^2+^-activated sinusoidal analysis cross-bridge kinetic model parameters by fiber type for 25 and 37°C. Mean ± SD, with each point representing an individual fiber. 2π*b* is the rate of myosin transition between the weakly and strongly-bound states (or rate of attachment), 2π*c* is the rate of myosin detachment from actin. 1-2: Rate of attachment and detachment with continuous y-axes to show the magnitude differences between fiber types and temperature. Where fiber type effects were observed, different letters above bars or brackets identify pair-wise differences (*p* < 0.05) between groups. Horizontal bars indicate significant (*p* < 0.1) interactions (temperature by fiber type). 3-4: Rate of attachment and detachment with different y-axes scales to visualize spread of data with fold-change plotted. Total number of fibers tested for each condition was MHC I (25°C: 147, 37°C: 98) and MHC IIA (25°C: 123, 37°C: 135).

Myosin attachment time, *t*_on_ calculated as (2π*c*)^-1^ (Palmer et al., 2007), became shorter with increased temperature for both fiber types (both *p* < 0.001), and was shorter in MHC IIA fibers than MHC I at both temperatures (MHC I: 5.5 ± 8.5 ms at 37°C vs. 40.6 ± 7.5 ms at 25°C, *p* < 0.001; MHC IIA: 3.4 ± 7.8 ms at 37°C vs. 16.3 ± 8.4 ms at 25°C, *p* = 0.0392). Additionally, in MHC I fibers, 2π*b* (8.7-fold) was more sensitive to temperature than 2π*c* (4.9-fold) with temperature, while MHC IIA fiber cross-bridge kinetics was not (both 4.0-fold). These alterations to cross-bridge kinetics with temperature can be further quantified by calculation of the Q_10_ temperature coefficient. Q_10_ values for 2π*b* and 2π*c* were greater for slow-contracting MHC I fibers compared to fast-contracting MHC IIA fibers, leading to a slow-to-fast Q_10_ ratio >1.0 (Table 4). Overall, these results show that higher temperatures greatly increased cross-bridge kinetics, with the largest changes in the slow-contracting MHC I fibers.

**Table 4.** Summary of Q_10_, the increase in a rate with an increase in temperature by 10°C, values from the current study compared to other literature.

| Study | Measurement | Temperature (°C) | Species | Slow Fibers | Fast Fibers | Slow-to-Fast Ratio |
| --- | --- | --- | --- | --- | --- | --- |
| Current study | $2\pi b$ (s<sup>-1</sup>) | 25-37 | Human | 6.34 | 3.73 | 1.7 |
| Wang & Kawai 2001 | $2\pi b$ (s<sup>-1</sup>) | 25-37 | Rabbit | 7.36 | 2.51 | 2.9 |
| Fenwick et al., 2021 | $2\pi b$ (s<sup>-1</sup>) | 17-27 | Mouse | 2.2 | | |
| Current study | $2\pi c$ (s<sup>-1</sup>) | 25-37 | Human | 4.93 | 3.66 | 1.3 |
| Wang & Kawai 2001 | $2\pi c$ (s<sup>-1</sup>) | 25-37 | Rabbit | 3.65 | 1.90 | 1.9 |
| Fenwick et al., 2021 | $2\pi c$ (s<sup>-1</sup>) | 17-27 | Mouse | 1.3 | | |
| Debold et al., 2004 | $V_{\max}$ (FL/s) ^ | 15-30 | Rat | 3.17 | 1.75 | 1.8 |
| Sundberg et al., 2018 | $V_{\max}$ (FL/s) ^ | 15-30 | Human | 3.20 | | |
| He et al., 2000 | $V_{\max}$ (FL/s) | 12-20 | Human | 3.85 | 3.46 | 1.1 |
| Knuth et al., 2006 | $V_0$ (FL/s) ^ | 15-30 | Rat | 4.20 | 3.07 | 1.4 |
| Bottinelli et al., 1996 | $V_0$ (FL/s) | 12-20 | Human | 6.98 | 4.98 | 1.4 |
| He et al., 2000 | Fiber ATPase (mM/s) | 12-20 | Human | 2.20 | 2.09 | 1.1 |

Fast = fast-contracting fibers, Slow = slow-contracting fibers, FL = fiber length, $V_{\max}$ = maximum shortening velocity, $V_0$ = unloaded shortening velocity, ^ indicates $Q_{10}$ calculated based upon values in other publications

Parameters *B* and *C* increased with temperature (*p* < 0.001) in both fiber types (Figures 4.1 and 4.2), indicating that 37°C results in an increase in the number of strongly-bound myosin-actin cross-bridges and/or their stiffness compared to 25°C. A significant interaction trend (*p* = 0.100) showed that the increase in *B* was more pronounced in MHC I fibers (86%) compared to MHC IIA fibers (34%); however, no interaction occurred for *C* as the increase did not differ between fiber types (58 and 70%, *p* = 0.268). In both MHC I and IIA fibers, *A* was unchanged with temperature (Figure 4.3, *p* = 0.788), while *k* was reduced (Figure 4.4, *p* < 0.001). Over the range of frequencies examined, the changes in *A* and *k* with higher temperature altered the elastic and viscous moduli (Figure 2.1-2.4) by 2 ± 4% (*p* = 0.403) and -14 ± 3% (*p* < 0.001)for MHC I fibers and by -14 ± 3% (*p* < 0.001) and -35 ± 2% (*p* < 0.001) for MHC IIA fibers, meaning that stiffness decreased in MHC IIA fibers and viscosity decreased in MHC I and IIA fibers at 37°C. At both 37°C and 25°C, MHC IIA fibers had larger *B* and *C* parameters (both *p* < 0.001) and larger *A* and smaller *k* (both p < 0.001) than MHC I fibers.

**Figure 4.**
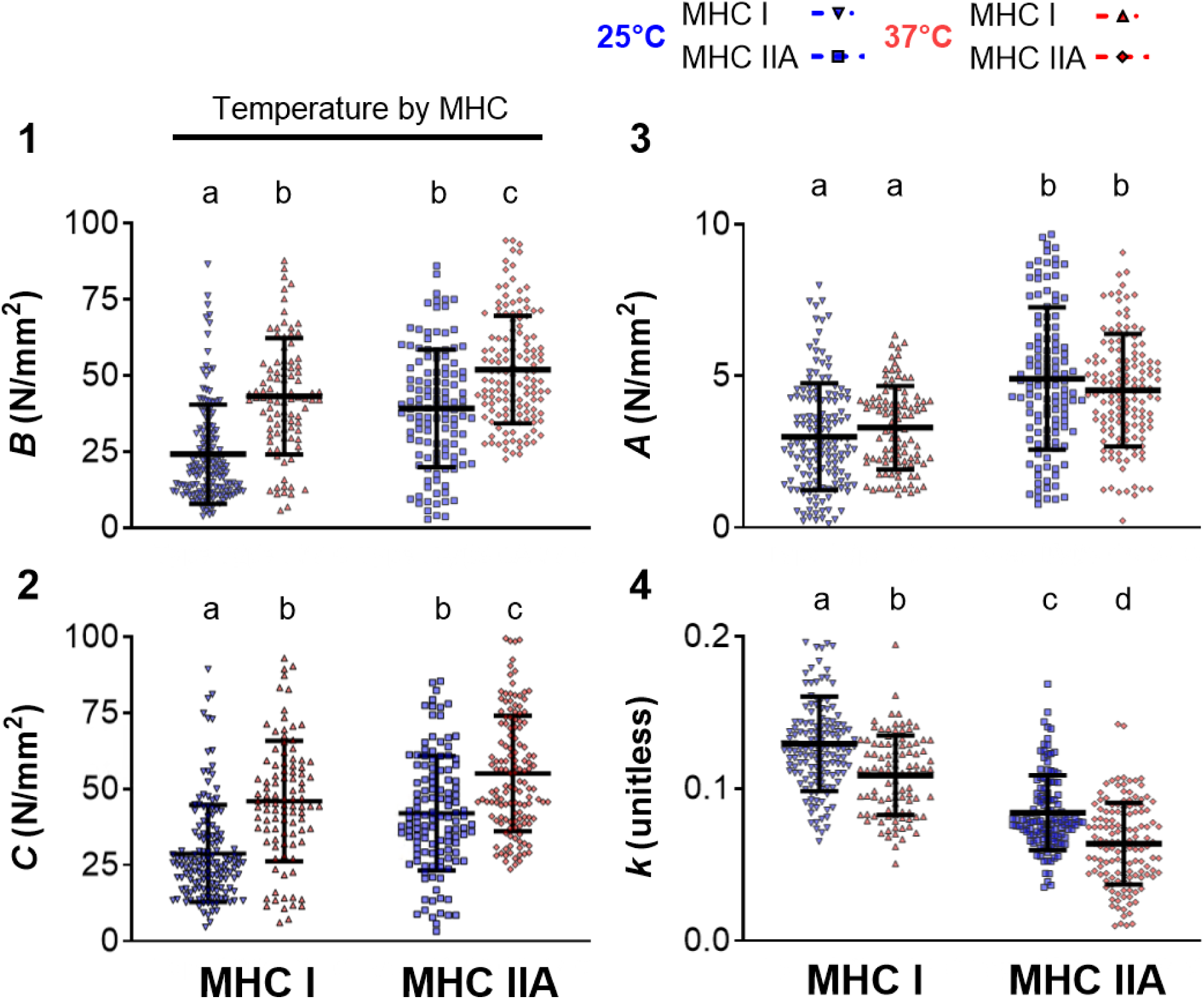
Single skeletal muscle fiber maximal Ca^2+^-activated sinusoidal analysis cross-bridge mechanical model parameters by fiber type for 25 and 37°C. Mean ± SD, with each point representing an individual fiber. 1-2: Parameters *B* and *C* are proportional to the number of myosin heads strongly-bound to actin and/or cross-bridge stiffness. 3-4: *A* and *k* represent the underlying lattice structure and attached myosin heads. Where fiber type effects were observed, different letters above bars or brackets identify pair-wise differences (*p* < 0.05) between groups. Horizontal bars indicate significant (*p* < 0.05) interactions (temperature by fiber type). Total number of fibers tested for each condition was MHC I (25°C: 147, 37°C: 98) and MHC IIA (25°C: 123, 37°C: 135).

### Effects of Temperature on Force Production Across Fiber Size

Because prior work from our laboratory and others have noted that specific tension declines with increasing CSA at various temperatures (5-25°C) (Elzinga *et al*., 1989; Gilliver *et al*., 2009; Miller *et al*., 2015), we evaluated whether this phenomenon occurs at 37°C. At 37°C, no relationship between specific tension and CSA existed, as indicated by a slope not different from zero in MHC I (Figure 5.1) and MHC IIA (Figure 5.2) fibers (Table 2). These results show that at physiological temperature the relationship between force and size produces consistent specific tensions, which is an underlying assumption when normalizing single fiber force to size and only examining specific tension values instead of plotting force per fiber size. In contrast, at 25°C, specific tension declined with CSA (Figure 5.1 and 5.2), as found previously at this temperature in fibers from young adults (Miller *et al*., 2015).

**Figure 5.**
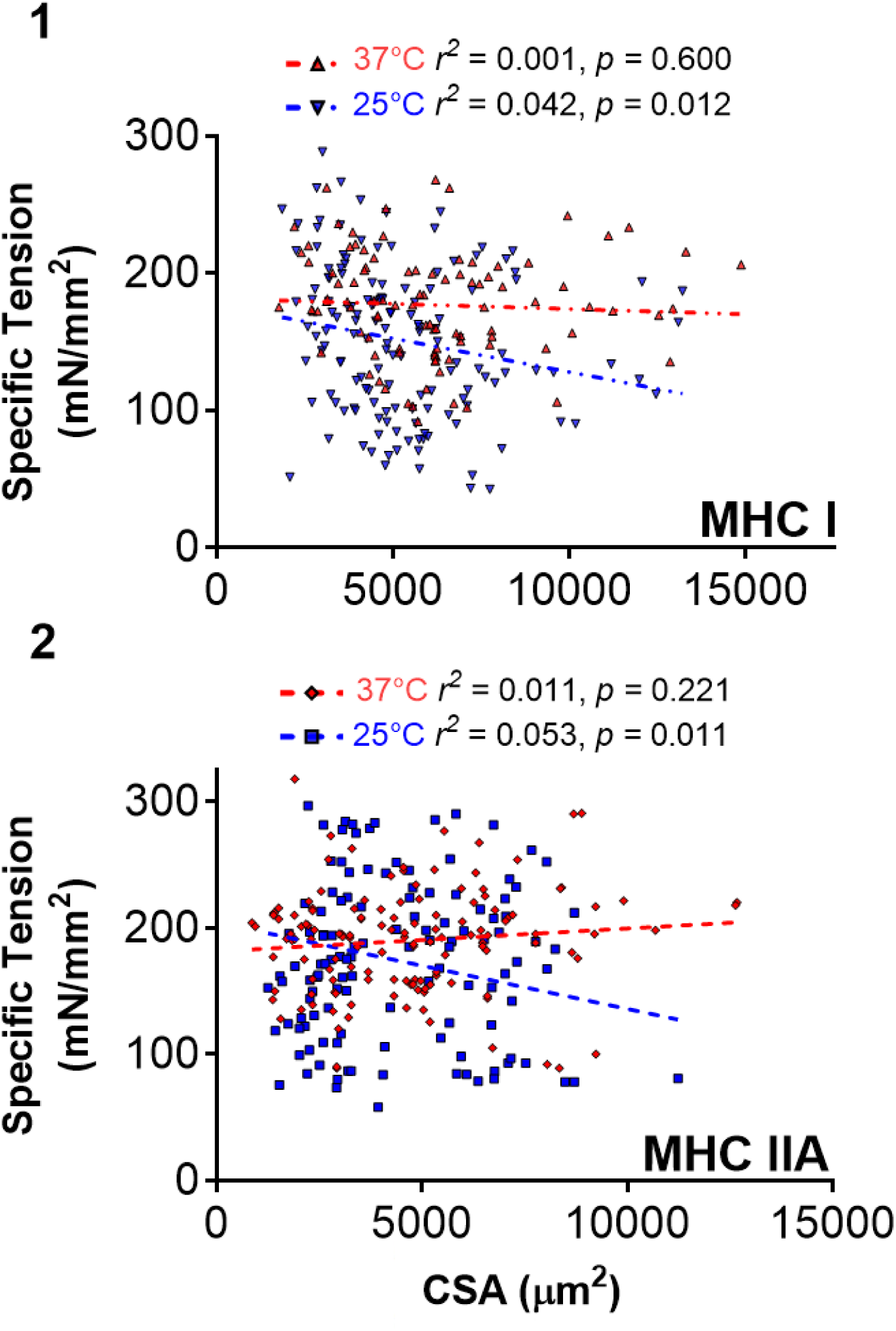
Relationship of isometric specific tension to CSA by fiber type for 25 and 37°C. Each point represents individual fibers with lines indicating linear regressions conducted for each fiber type (MHC I or IIA) and for each condition 25 or 37°C. Specific tension and CSA were negatively correlated at 25°C for both fiber types (p ≤ 0.05), whereas no correlation was observed at 37°C (p > 0.05). Slopes and intercepts for lines are as indicated: MHC I 25°C y = - 4.89E^-03^x + 176.81; MHC I 37°C y = 7.62E^-04^x + 181.54; MHC IIA 25°C y = -6.82E^-03^x + 204.06; MHC IIA 37°C y = 1.83E^-03^x + 181.27. Total number of fibers tested for each condition was MHC I (25°C: 147, 37°C: 98) and MHC IIA (25°C: 123, 37°C: 135).

### Effects of Temperature on Cross-Bridge Kinetics Across Fiber Size

To examine the effects of fiber size on myofilament mechanics, we performed correlations between CSA and the six curve-fit parameters. No change in cross-bridge kinetics, 2π*b* or 2π*c*, occurred across fiber size (Figure 6, Table 2) for either fiber type, as their slopes were not different from zero at either temperature. These findings indicate that while substantial increases in cross-bridge kinetics occur in both MHC I and IIA fibers when transitioning to 37°C, fiber size alone does not lead to alterations in cross-bridge kinetics at either temperature and, thus, does not likely explain temperature changes in the specific tension versus CSA response (Figure 5). These findings align with prior work from our laboratory at 25°C in that cross-bridge kinetics have no relationship with CSA in fibers from young adults (Miller *et al*., 2015).

**Figure 6.**
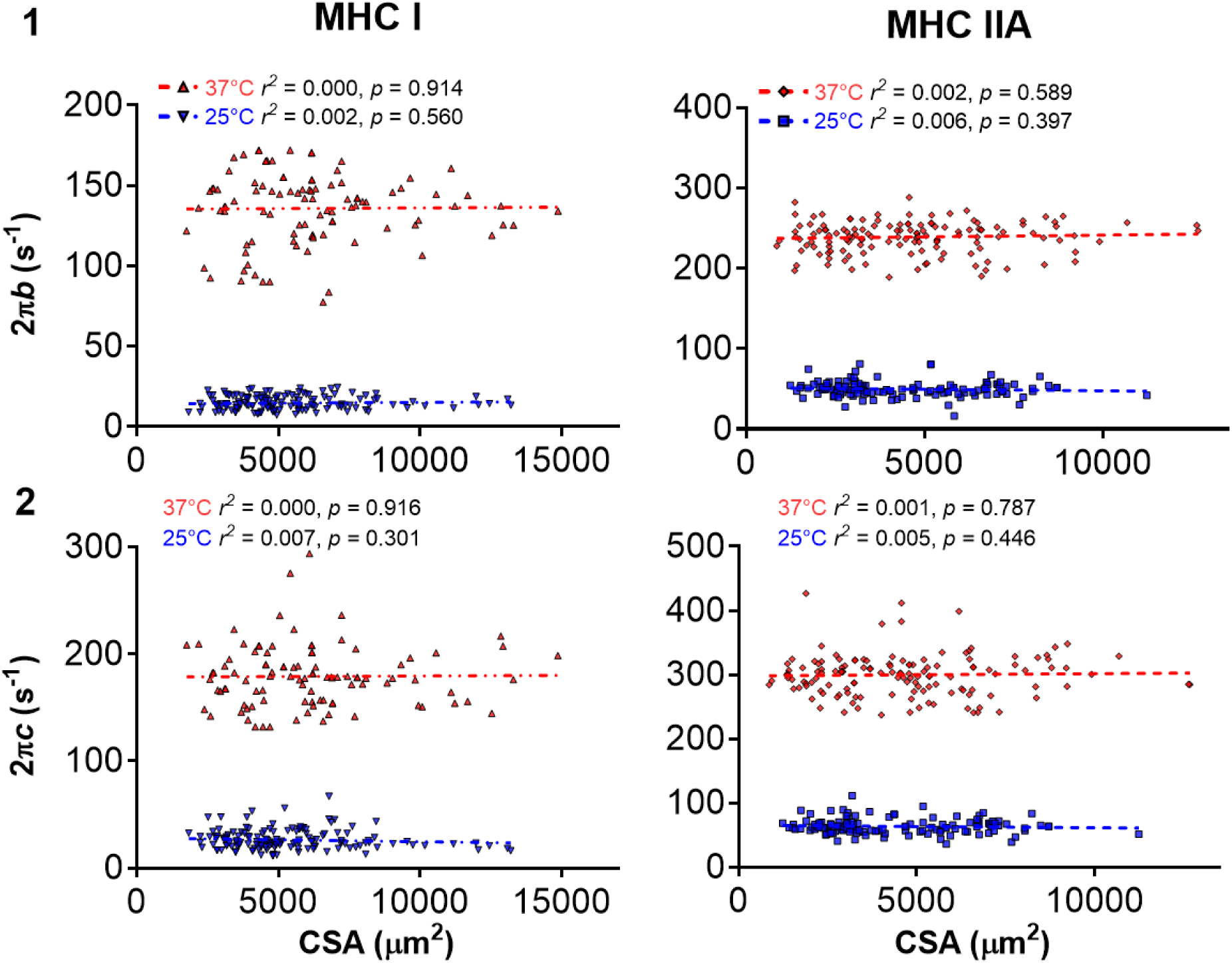
Relationship of sinusoidal analysis model parameters 2π*b* (1) and 2π*c* (2) to CSA for by fiber type for 25 and 37°C. Each point represents individual fibers with lines indicating linear regressions conducted for each fiber type (MHC I or IIA) and for each condition 25 or 37°C. Myosin-actin cross-bridge kinetics did not change with CSA for either temperature or fiber type, as their slopes were not different from zero (*p* > 0.05). Slopes and intercepts for lines are as indicated: 2π*b*: MHC I 25°C y = 9.04E^-05^x + 14.28; MHC I 37°C y = 8.88E^-05^x + 135.29; MHC IIA 25°C y = -3.58E^-04^x + 51.32; MHC IIA 37°C y = 4.47E^-04^x + 236.69. 2π*c*: MHC I 25°C y = -3.56E^-04^x + 28.36; MHC I 37°C y = 1.13E^-04^x + 178.44; MHC IIA 25°C y = -3.92E^-04^x + 65.42; MHC IIA 37°C y = 3.46E^-04^x + 298.34. Total number of fibers tested for each condition was MHC I (25°C: 147, 37°C: 98) and MHC IIA (25°C: 123, 37°C: 135).

### Effects of Temperature on Strongly-Bound Cross-Bridges Across Fiber Size

Parameters typically normalized by dividing by CSA (*A*, *B* and *C*) were unnormalized (*A*_CSA_, *B*_CSA_, and *C*_CSA_) in order to remove their direct relationship to fiber size and thus represent absolute magnitudes. *A*_CSA_ increased with fiber size at both temperatures, while *k* was not related to fiber size at 37°C or for MHC IIA at 25°C, but decreased with size in MHC I fibers at 25°C (Table 2). *B*_CSA_ and *C*_CSA_ were the parameters most highly correlated (*r*) with fiber size (Table 2) and had similar relationships to CSA, so only *C*_CSA_ data are plotted. C_CSA_ versus CSA plots indicated that MHC I and IIA fibers have a greater number of strongly-bound cross-bridges or cross-bridge stiffness at 37°C compared to 25°C as indicated by a steeper slope (Figure 7.1). Force versus C_CSA_ plots showed that both MHC I and IIA fibers produce similar amounts of force for any given number or stiffness of strongly-bound cross-bridges (Figure 7.2). Force and C_CSA_ increase similarly with fiber size, as shown by their ratio (Force / C_CSA_) being constant over the range of CSAs as their slopes were not different from zero (Figure 8.1). Force / C_CSA_ was similar in MHC I and IIA fibers at 25 and 37°C. The variability when collected at 37°C is substantially less than at 25°C (Figure 8.2), meaning these measurements taken at physiological temperatures have more consistent results.

**Figure 7.**
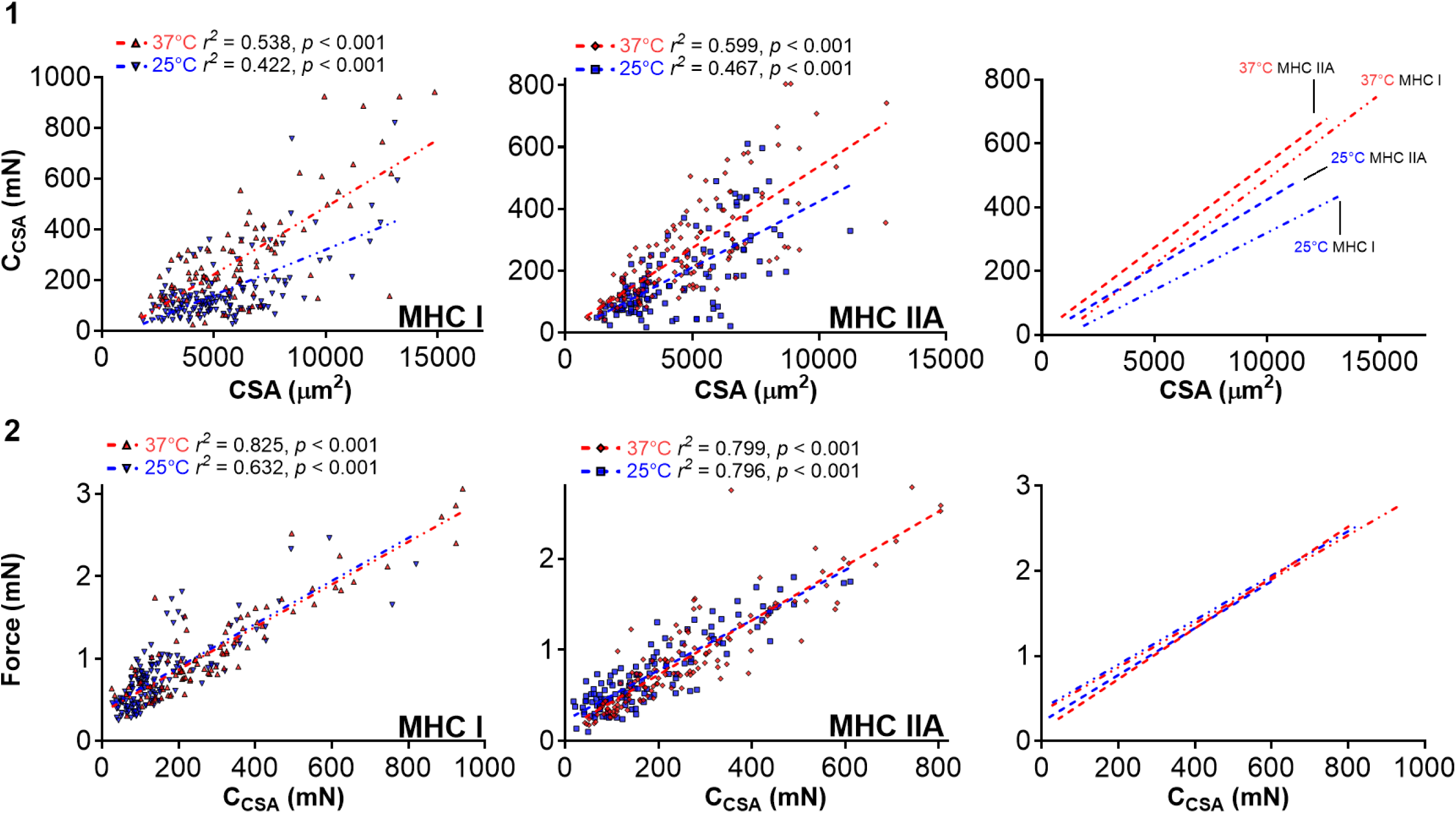
Relationship of strongly-bound cross-bridges and force or CSA for by fiber type for 25 and 37°C. Each point represents individual fibers with lines indicating linear regressions conducted for each fiber type (MHC I or IIA), and for each condition 25 or 37°C. 1: MHC I and IIA fibers had stronger relationships (r^2^ value in figure or r value in Table 2) between absolute magnitude of cross-bridge number or stiffness and CSA. ANCOVA analysis identified slopes between temperatures for each fiber type MHC I and IIA were significantly different (*p* < 0.001) but not intercepts (MHC I: *p* = 0.608; MHC IIA: *p* = 0.758). 2: Force and strongly-bound cross-bridges have a strong relationship and each fiber type produced similar force for a given cross-bridge number and/or stiffness. ANCOVA analysis identified slopes (MHC I: *p* = 0.0862; MHC IIA: *p* = 0.0994) and intercepts (MHC I: *p* = 0.118; MHC IIA: *p* = 0.683) were not significantly different between temperatures. Slopes and intercepts for lines are as indicated: CSA-C_CSA_: MHC I 25°C y = 0.036x - 22.76; MHC I 37°C y = 0.053x - 43.01; MHC IIA 25°C y = 0.043x + 1.38; MHC IIA 37°C y = 0.053x + 9.54. Force-C_CSA_: MHC I 25°C y = 2.61E^-03^x + 0.37; MHC I 37°C y = 2.58E^-03^x + 0.35; MHC IIA 25°C y = 2.75E^-03^x + 0.22; MHC IIA 37°C y = 2.98E^-03^x + 0.13. Total number of fibers tested for each condition was MHC I (25°C: 147, 37°C: 98) and MHC IIA (25°C: 123, 37°C: 135).

**Figure 8.**
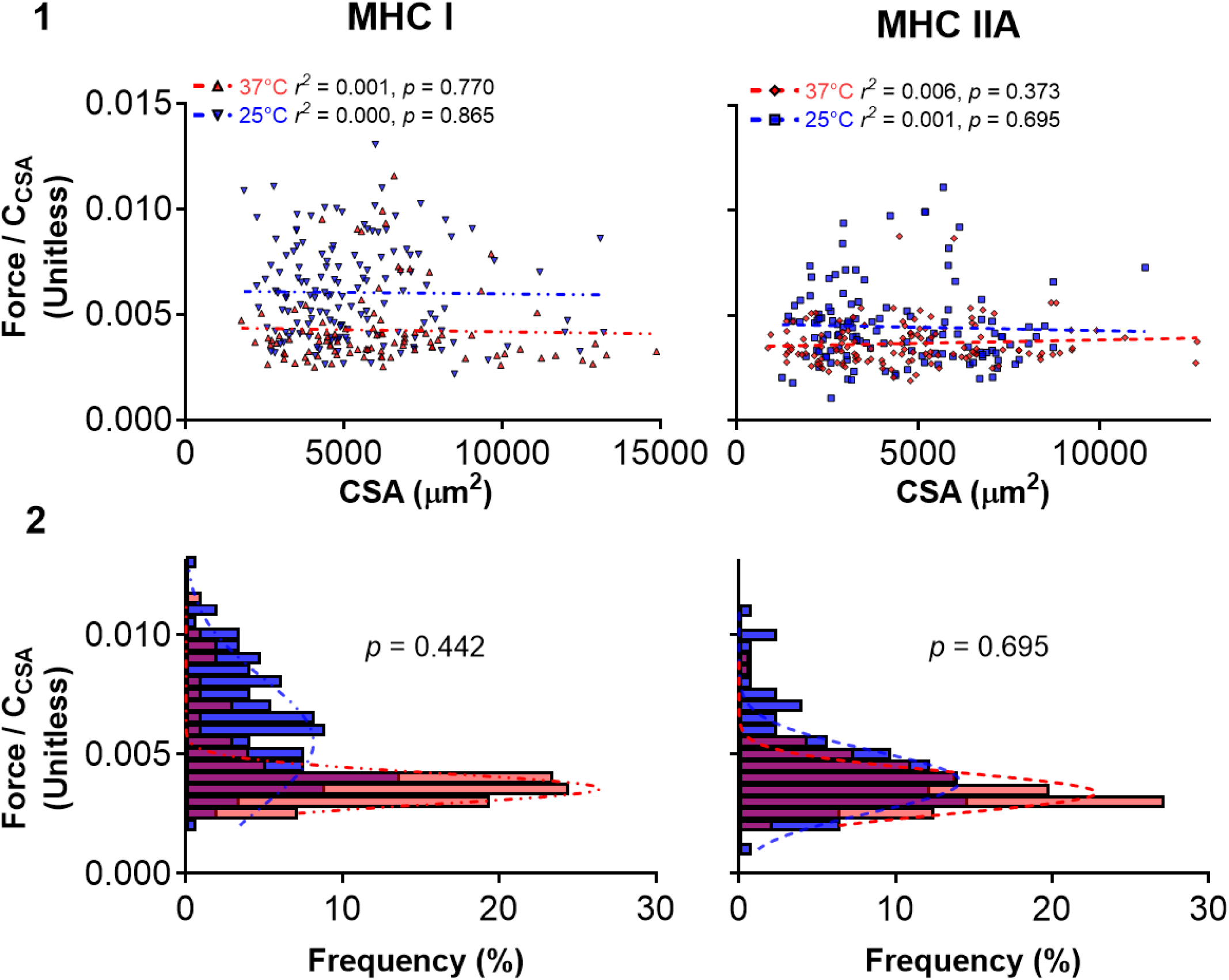
Relationship of force/C_CSA_ and CSA and frequency distribution by fiber type for 25 and 37°C. 1: Each point represents individual fibers with lines indicating linear regressions conducted for each fiber type (MHC I or IIA), and for each condition 25 or 37°C. MHC I and IIA fibers had no relationship between force/C_CSA_ and CSA for 25 or 37°C. Slopes and intercepts for regression lines are as follows: MHC I 25°C y = -1.41E^-08^x + 0.006; MHC I 37°C y = -1.96E^-08^x + 0.004; MHC IIA 25°C y = -3.25E^-08^x + 0.004; MHC IIA 37°C y = -3.28E^-08^x + 0.003. 2: Frequency distribution of force/C_CSA_ data for each condition. Total number of fibers tested for each condition was MHC I (25°C: 147, 37°C: 98) and MHC IIA (25°C: 123, 37°C: 135).

## DISCUSSION

The results of this work from skeletal muscle fibers in older males and females (66-76 years) advance our understanding of human single fiber and molecular-level contractile function at body temperature (37°C). Compared with 25°C, contractile function at 37°C was more similar between MHC I and IIA fibers, as evidenced by the lack of difference in specific tension (Figure 1.1), a reduced gap in cross-bridge kinetics (Figure 3), and overlap in oscillatory work and power output (Figure 2.5-2.6) between fiber types. This similarity was established by a more pronounced response to the increase from 25°C to 37°C in MHC I fibers, especially in terms of myosin-actin cross-bridge kinetics and number or stiffness of strongly-bound cross-bridges.

Notably, cross-bridge kinetics were the variables most affected by increased temperature in both fiber types, increasing by 4.7-9.4-fold. The relationship between fiber force and size was stronger at 37°C in both fiber types with r^2^ values ranging from 0.80-0.82 compared to 0.51-0.52 at 25°C, meaning that CSA explains a greater proportion of force at body temperature. This improved relationship between fiber force and size at 37°C, which was likely due to the increased number of strongly-bound cross-bridges at 37°C, led to specific tension being independent of size, a desired outcome for the measure of normalized force. Overall, these results indicate that performing experiments at body temperature provides unique insights into single fiber and molecular level contractile performance, which may lead to a better understanding of whole muscle contractility *in vivo*.

Our current findings indicate that specific tension is not different between MHC isoforms at 37°C. In contrast, our data collected at 25°C agree with many studies completed at sub-physiological temperatures (≤30°C) that indicated fast-contracting fibers produce more force per muscle size than slow-contracting fibers from human muscle (Bottinelli *et al*., 1996; D’Antona *et al*., 2003; Choi & Widrick, 2010; Krivickas *et al*., 2011; Hvid *et al*., 2011; Miller *et al*., 2015). In addition to specific tension, studies at lower temperatures (≤30°C) have shown that fast-contracting fibers produce significantly more power (force x velocity) than slow-contracting fibers, mostly because fast-contracting fibers produce higher contractile velocities (Bottinelli *et al*., 1991, 1996). Our current results indicate power production should be comparable between fiber types, as specific tension was not different and the rate of myosin attachment and detachment become more similar between fiber types at 37°C.

We demonstrated that cross-bridge kinetics in MHC I fibers are more temperature-sensitive than MHC IIA fibers, increasing 6.7-9.4-fold in MHC I fibers and only 4.7-4.8-fold in MHC IIA fibers (Figure 3). Prior studies using muscle from rabbits and rats showed that the Q_10_, especially the rate of cross-bridge attachment (Wang & Kawai, 2001), is higher in slow-contracting fibers than fast-contracting at temperatures ranging from 15-37°C (Wang & Kawai, 2001; Debold *et al*., 2004; Knuth *et al*., 2006) (Table 4). In human skeletal muscle studied between 12-20°C, findings have been equivocal, with some studies indicating a higher Q_10_ in slow-contracting fibers (Bottinelli *et al*., 1996), while others found no difference (He *et al*., 2000). The greater Q_10_ values for 2π*b* than 2π*c* noted in the current study and in previous work (Wang & Kawai, 2001) indicate that more myosin heads are transitioning into the strongly-bound state than detaching from actin in slow-contracting fibers at higher temperatures, which agrees with the increased specific tension in these fibers at 37°C compared to 25°C. With the assumption that contractile velocity is proportional to the cross-bridge unitary displacement (d_uni_, nm) divided by *t*_on_ (ms) (Tyska & Warshaw, 2002; Miller & Toth, 2013), which is a detachment-limited model of velocity (Huxley, 1990), a more similar *t*_on_ should lead to more similar velocities. Contractile velocity has been linked to myosin attachment rate as well (Hooft *et al*., 2007; Walcott *et al*., 2012), suggesting that our finding of a more sensitive attachment rate in MHC I than MHC IIA fibers could indicate more comparable contractile velocities at physiological temperatures. Overall, our results at 37°C indicate no difference between MHC I and IIA fibers in specific tension (Figure 1.1) and more similar myosin-actin cross-bridge kinetics than at 25°C (Figure 3), which may lead to more similar contractile velocities, as found at 35°C for motility speed between the various human muscle fiber types (Lionikas *et al*., 2006).

Prior studies seeking to understand the mechanisms of increased force generation at higher temperatures have generally indicated that increased force per cross-bridge (Ford *et al*., 1977; Kuhn *et al*., 1979; Goldman *et al*., 1987; Dantzig *et al*., 1992; He *et al*., 2000; Piazzesi *et al*., 2003; Decostre *et al*., 2005; Ranatunga, 2010, 2018; Brenner *et al*., 2012), an increase in cross-bridge number (Wang & Kawai, 2001; Kawai, 2003), or a combination of both (He *et al*., 2000; Brenner *et al*., 2012) are responsible for greater force at higher temperatures. Past work has identified that, with increasing temperatures, there is a relatively greater increase in isometric force compared to fiber stiffness (Ford *et al*., 1977; Kuhn *et al*., 1979; Goldman *et al*., 1987; Dantzig *et al*., 1992; He *et al*., 2000; Piazzesi *et al*., 2003; Decostre *et al*., 2005; Ranatunga, 2010, 2018; Brenner *et al*., 2012). These findings have been interpreted as an increase in force per cross-bridge, with the assumption that the individual cross-bridge does not contribute substantially to fiber stiffness. The prevailing thought behind an increased force per cross-bridge is a temperature-induced increase in cross-bridge strain (Linari *et al*., 2007; Brenner *et al*., 2012). However, others have argued that greater force at higher temperatures is solely due to increased cross-bridge number (Wang & Kawai, 1997; Kawai, 2003). This latter interpretation is based on the observation that a series elastic model can adequately describe the change in force with minimal change in stiffness from increased temperature due to the compliance of the thick and thin filaments (Wakabayashi *et al*., 1994; Kojima *et al*., 1994). This model indicates that I-band stiffness limits the increase in stiffness of single fibers, and the relatively smaller increase in fiber stiffness is due to the increase in cross-bridge number (Wang & Kawai, 2001). Notably, many of these studies have been conducted on animal tissues and may not translate to human skeletal muscle, as humans may rely on both increased cross-bridge number and force per cross-bridge (He *et al*., 2000; Brenner *et al*., 2012). These prior findings suggest that translating from lower temperatures or animal tissue to *in vivo* conditions in humans may have limitations, highlighting the critical importance of studying human skeletal muscle tissue at 37°C.

Our C_CSA_ findings indicate that the increase in specific tension with temperature in both fiber types (Figure 1) is primarily due to an increase in the number of strongly-bound cross-bridges and not changes in force per cross-bridge or cross-bridge stiffness. If force per cross-bridge alone was increased with temperature, then fibers at 37°C would produce a greater force for the same C_CSA_ at 25°C. In other words, the force versus C_CSA_ curve at 37°C should have greater force values across the range of C_CSA_s compared to 25°C; however, we observed the same force values across the range of C_CSA_s for both temperatures (Figure 7.2). Separating out whether C_CSA_ represents the number of strongly-bound cross-bridges or cross-bridge stiffness is more difficult. As fibers with larger CSAs should have more strongly-bound cross-bridges, but not necessarily greater cross-bridge stiffness, our finding that force and C_CSA_ increase similarly with fiber size, as shown by their ratio (Force / C_CSA_) being constant over the range of CSAs (Figure 8.1), strongly suggests C_CSA_ is a measure of strongly-bound cross-bridges. Additionally, as the A-process represents the underlying lattice structure as well as the myosin heads attached in series (Mulieri *et al*., 2002; Palmer *et al*., 2004), our finding of no change in A (Figure 4.3) and a decrease in k (Figure 4.4), which leads to elastic modulus being unchanged in MHC I fibers and decreased in MHC IIA fibers also indicates that cross-bridge stiffness is not significantly increasing to produce more force.

A potential mechanism to explain the increase in strong-binding with higher temperatures is that a population of myosin heads at rest may move from a disordered refractory-like state that is unable to bind myosin at a lower temperature (10°C) to an ordered state able to bind myosin at higher temperatures (35°C) upon activation of the muscle (Caremani *et al*., 2019). However, those results from intact mouse extensor digitorum longus (Caremani *et al*., 2019) were not observed in skinned psoas rabbit fibers at temperatures ranging from 5-20°C (Linari *et al*., 2007). The discrepancies between these prior studies are potentially due to lattice swelling, as at higher temperatures (35-37°C) intact and skinned rabbit psoas skeletal and rat cardiac muscle at rest show similar thick filament structure if the skinned myofilament lattice is compressed with Dextran (Caremani *et al*., 2021; Ovejero *et al*., 2022).

Despite not compressing the myofilament lattice here, our results in skinned human skeletal muscle appear consistent with the results from intact mice in that more myosin heads are available to bind actin at higher temperatures. Additionally, our results that higher temperature increases the number of strongly-bound cross-bridges agree well with previous studies in animals using similar techniques (Wang & Kawai, 1997; Kawai, 2003), but only partially agree with previous human studies at lower temperatures that indicate temperature increases both cross-bridge number and force per cross-bridge (He *et al*., 2000; Brenner *et al*., 2012). Future experiments using different techniques in human tissue at body temperatures are needed to verify or contradict our findings. Additionally, studies examining different ages or health status may be needed as our work focused on muscle fibers from older adults, which perform differently in various aspects compared with fibers from young adults (Frontera *et al*., 2000; D’Antona *et al*., 2003; Hvid *et al*., 2011; Miller & Toth, 2013).

As force production is related to the number of strongly-bound cross-bridges, force generated per cross-bridge, and myofilament stiffness (Tanner *et al*., 2007), we examined parameters *A* and *k*, which relate to the viscoelastic properties of the myofilament lattice structure and attached myosin heads in series (Mulieri *et al*., 2002; Palmer *et al*., 2004). Our data, collected while cross-bridges were cycling, indicate greater myofilament compliance at higher temperatures, in agreement with our previous work under relaxed conditions, where cross-bridges are unattached (Palmer *et al*., 2013). Studies using rigor (no ATP) to investigate fiber function, where >94% of myosin heads that can bind will be in a strongly-bound state (Cooke & Franks, 1980; Lovell *et al*., 1981), show lower specific tension at higher temperatures (Goldman *et al*., 1987). Assuming at higher temperatures the number of strongly bound cross-bridges will be similar or greater due to more heads being available to bind actin (Caremani *et al*., 2019) and force generated per cross-bridge will be similar, these results also suggest higher temperatures cause a decrease in myofilament stiffness. Overall, we interpret our results to indicate that, in human skeletal muscle, an increase to body temperature induces a greater number of strongly-bound cross-bridges with increased myofilament compliance.

The validity of using average specific tension to generalize findings has been called into question because at lower temperatures (5-25°C) specific tension declines as fiber size increases, as found here (Figure 5), in our previous work (Miller *et al*., 2015), and in the work of others (Elzinga et al., 1989; Gilliver et al., 2009; Meijer et al., 2015; Jeon et al., 2019). Several groups have proposed mechanisms for the loss of specific tension with greater cross-sectional area, including altered cross-bridge kinetics with increasing fiber size from ADP or phosphate accumulation (Elzinga *et al*., 1989; Gilliver *et al*., 2009), errors in measuring fiber size (Smith & Herzog, 2023), and differences in myofilament density or myosin in larger fibers (Miller *et al*., 2015; Meijer *et al*., 2015). Our work in fibers from young (Miller *et al*., 2015) and older adults (Figure 6) show that cross-bridge kinetics, at least in terms of the rate of myosin transition between the weakly and strongly-bound states and the rate of myosin detachment, are similar across fiber size. As such, these data indicate that the suggested accumulation of small molecules, such as ADP or phosphate, due to diffusion limitations in larger fibers (Elzinga *et al*., 1989; Gilliver *et al*., 2009) is not occurring. Recent experiments also show that measuring single fiber diameters in horizontal and vertical directions and calculating skinned single fiber CSA as an ellipse results in ∼2% error after 30 minutes of skinning and <1% error after 60 minutes of skinning compared to actual single fiber CSA (Mebrahtu *et al*., 2024). Even when calculating skinned single fiber CSA from one vertical measurement and assuming the fiber was a circle resulted in <2% error after 60 minutes of skinning (Mebrahtu *et al*., 2024). Collectively, these results suggest that errors in fiber size measurements are most likely not the underlying reason for reductions in specific tension as CSA increases. In the current study, we skinned fibers for ∼60 minutes (muscle bundle for 30-40 minutes before fiber removal and then single fibers for an additional 30 minutes) and CSA was calculated as an ellipse after measuring diameters horizontally and vertically. Our results at 37°C indicate no relationship between specific tension and CSA (Figure 5), potentially due to a stronger force-CSA relationship at higher temperatures. For instance, CSA explained 80-82% of the variance in force at 37°C, which is likely due to a greater proportion of the available cross-bridges being bound, compared to 51-52% at 25°C (Figure 7.2). These findings strongly indicate that measuring single fiber performance at 37°C produces different results than at 25°C. However, this does not mean that measurements at lower temperatures are invalid, but future reports that include plots of the force versus CSA in addition to the typical plots of specific tension may enhance our understanding of the relationship between these two variables.

This work in tissue from healthy older adults suggests that MHC I and IIA fibers perform more similarly at physiological temperatures when observing force production, oscillatory work and power, and cross-bridge kinetics (Figures 1-3). *In vivo*, different fiber types (MHC I, IIA, IIX and hybrids I/IIA, IIA/IIX) have been thought to help muscle work under a wide range of force-velocity profiles to fine-tune muscle performance or to MHC isoform transition due to exercise or aging (Stephenson, 2001; Medler, 2019). Prior studies have shown that hybrid MHC isoforms have intermediate contractile properties for cross-bridge kinetics (Galler *et al*., 1994; Miller *et al*., 2015), unloaded and maximal shortening velocities (Bottinelli *et al*., 1994*a*, 1996), and myofibrillar ATPase (Bottinelli *et al*., 1994*b*). These findings suggest that hybrid fibers aid the whole-muscle’s ability to scale force and contractile velocity to the task demand (Stephenson, 2001; Medler, 2019). But a recent review has indicated that in healthy, physically active populations, the proportion of hybrid fibers is substantially lower compared to pure MHC I and IIA fibers (Medler, 2019). If functional properties of MHC I and IIA fibers are more similar in healthy-states at physiological temperatures, as reported here, the need for hybrid fibers to provide intermediate contractile function between pure fibers may be lessened. However, in states of disease or disuse, the abundance of hybrid or fast-contracting fibers increases, potentially out of necessity to allow for physical task demand completion (Medler, 2019). Our prior work in humans and animal models indicates that with age or disease, cross-bridge kinetics become slower across a range of fiber types (Miller *et al*., 2010, 2013; Toth *et al*., 2013; Momb *et al*., 2022, 2023*a*). Depending on the extent of the slowing of MHC I fibers and IIA fibers in these instances, these may provoke the transition to I/IIA or IIA/IIX fibers, suggesting a physiologic benefit for these fibers to overcome a functional deficits that may present from slowed kinetics.

In conclusion, these results highlight that single fiber performance depends significantly on the temperature studied, particularly in more temperature-sensitive, slower-contracting MHC I fibers. Human physiological temperature (37°C) leads to slow- and fast-contracting fibers acting more alike than different; producing comparable specific tensions, oscillatory work and power that overlap, and closing the gap on cross-bridge kinetic rates compared with a lower temperature (25°C). In both fiber types, fiber CSA explains a greater proportion of force at body temperature than at 25°C. This stronger relationship between fiber force and size at 37°C, which is likely due to the increased number and/or stiffness of strongly-bound cross-bridges, led to specific tension being independent of size, a desired outcome for the measure of normalized fiber force that is commonly not found at lower temperatures. Overall, these results indicate that performing experiments at 37°C provides unique insights into human single fiber and molecular-level contractile performance, which may lead to a better understanding of whole muscle contractility *in vivo*.

## ADDITIONAL INFORMATION

### Data availability statement

All data are represented within this manuscript.

### Competing interests

The authors declare that they have no competing interests.

### Author contributions

B.A.M and M.S.M. conceived and designed the research; B.A.M and S.R.C. acquired human skeletal muscle tissue; B.A.M. performed experiments; B.A.M. analyzed data; B.A.M, J.A.K, S.R.C., and M.S.M interpreted results of research; B.A.M., J.A.K and M.S.M prepared figures; B.A.M. and M.S.M. drafted manuscript; All authors contributed to the critical revision of the manuscript and approved the final version of the manuscript submitted for publication.

### Funding

This work was supported by the National Institutes of Health Grants R01 AG-047245 and R01 AG-058607.

## Acknowledgements

We thank the volunteers who dedicated their valuable time to these studies and the Center for Human Health and Performance in the Institute for Applied Life Sciences at the University of Massachusetts, Amherst.

## First author’s present address

30 Eastman Ln

Totman Building, Room 140E

Amherst, MA

01003

## Notes

### Competing Interest Statement

The authors have declared no competing interest.

## REFERENCES

Awinda PO, Bishaw Y, Watanabe M, Guglin MA, Campbell KS & Tanner BCW (2020). Effects of mavacamten on Ca2+ sensitivity of contraction as sarcomere length varied in human myocardium. Br J Pharmacol 177, 5609–5621.

Bergh U & Ekblom B (1979). Influence of muscle temperature on maximal muscle strength and power output in human skeletal muscles. Acta Physiol Scand 107, 33–37.

Bottinelli R, Betto R, Schiaffino S & Reggiani C (1994a). Maximum shortening velocity and coexistence of myosin heavy chain isoforms in single skinned fast fibres of rat skeletal muscle. J Muscle Res Cell Motil 15, 413–419.

Bottinelli R, Canepari M, Pellegrino MA & Reggiani C (1996). Force-velocity properties of human skeletal muscle fibres: myosin heavy chain isoform and temperature dependence. J Physiol 495, 573–586.

Bottinelli R, Canepari M, Reggiani C & Stienent GJM (1994b). Myofibrillar ATPase activity during isometric contraction and isomyosin composition in rat single skinned muscle fibres. J Physiol 481, 663–675.

Bottinelli R, Schiaffino S & Reggiani C (1991). Force-velocity relations and myosin heavy chain isoform compositions of skinned fibres from rat skeletal muscle. J Physiol 437, 655–672.

Brenner B (2006). The stroke size of myosins: A reevaluation. J Muscle Res Cell Motil 27, 173– 187.

Brenner B, Hahn N, Hanke E, Matinmehr F, Scholz T, Steffen W & Kraft T (2012). Mechanical and kinetic properties of β-cardiac/slow skeletal muscle myosin. J Muscle Res Cell Motil 33, 403–417.

Callahan DM, Miller MS, Sweeny AP, Tourville TW, Slauterbeck JR, Savage PD, Maugan DW, Ades PA, Beynnon BD & Toth MJ (2014). Muscle disuse alters skeletal muscle contractile function at the molecular and cellular levels in older adult humans in a sex-specific manner. J Physiol 592, 4555–4573.

Capitanio M, Canepari M, Cacciafesta P, Lombardi V, Cicchi R, Maffei M, Pavone FS & Bottinelli R (2006). Two independent mechanical events in the interaction cycle of skeletal muscle myosin with actin. Proc Natl Acad Sci U S A 103, 87.

Caremani M, Brunello E, Linari M, Fusi L, Irving TC, Gore D, Piazzesi G, Irving M, Lombardi V & Reconditi M (2019). Low temperature traps myosin motors of mammalian muscle in a refractory state that prevents activation. J Gen Physiol 151, 1272.

Caremani M, Fusi L, Linari M, Reconditi M, Piazzesi G, Irving TC, Narayanan T, Irving M, Lombardi V & Brunello E (2021). Dependence of thick filament structure in relaxed mammalian skeletal muscle on temperature and interfilament spacing. J Gen Physiol 153, e202012713.

Chase PB & Kushmerick MJ (1988). Effects of pH on contraction of rabbit fast and slow skeletal muscle fibers. Biophys J 53, 935–946.

Chen X, Sanchez GN, Schnitzer MJ & Delp SL (2016). Changes in sarcomere lengths of the human vastus lateralis muscle with knee flexion measured using in vivo microendoscopy. J Biomech 49, 2989–2994.

Choi SJ & Widrick JJ (2010). Calcium-activated force of human muscle fibers following a standardized eccentric contraction. Am J Physiol Cell Physiol 299, 1409–1417.

Cooke R & Franks K (1980). All Myosin Heads form Bonds with Actin in Rigor Rabbit Skeletal Muscle. Biochemistry 19, 2265–2269.

Coupland ME, Puchert E & Ranatunga KW (2001). Temperature dependance of active tension in mammalian (rabbit psoas) muscle fibres: Effect of inorganic phosphate. J Physiol 536, 879–891.

Coupland ME & Ranatunga KW (2003). Force generation induced by rapid temperature jumps in intact mammalian (rat) skeletal muscle fibres. J Physiol 548, 439–449.

D’Antona G, Lanfranconi F, Pellegrino MA, Brocca L, Adami R, Rossi R, Moro G, Miotti D, Canepari M & Bottinelli R (2006). Skeletal muscle hypertrophy and structure and function of skeletal muscle fibres in male body builders. J Physiol 570, 611–627.

D’Antona G, Pellegrino MA, Adami R, Rossi R, Naccari Carlizzi C, Canepari M, Saltin B & Bottinelli R (2003). The effect of ageing and immobilization on structure and function of human skeletal muscle fibres. J Physiol 552, 499–511.

Dantzig JA, Goldman YE, Millar NC, Lacktis J & Homsher E (1992). Reversal of the cross-bridge force-generating transition by photogeneration of phosphate in rabbit psoas muscle fibres. J Physiol 451, 247–278.

Debold EP, Dave H & Fitts RH (2004). Fiber type and temperature dependence of inorganic phosphate: Implications for fatigue. Am J Physiol Cell Physiol 287, 673–681.

Decostre V, Bianco P, Lombardi V & Piazzesi G (2005). Effect of temperature on the working stroke of muscle myosin. Proc Natl Acad Sci U S A 102, 13927–13932.

Elzinga G, Stienen GJM & Wilson MGA (1989). Isometric force production before and after chemical skinning in isolated muscle fibres of the frog rana temporaria. J Physiol 410, 171– 185.

Flouris AD, Webb P, Kenny GP & Kenny GP (2015). Noninvasive assessment of muscle temperature during rest, exercise, and postexercise recovery in different environments. J Appl Physiol 118, 1310–1320.

Ford LE, Huxley AF & Simmons RM (1977). Tension responses to sudden length change in stimulated frog muscle fibres near slack length. J Physiol 269, 441.

Fradkin AJ, Zazryn TR & Smoliga JM (2010). Effects of warming-up on physical performance: A systematic review with meta-analysis. J Strength Cond Res 24, 140–148.

Frontera WR, Suh D, Krivickas LS, Hughes VA, Goldstein R & Roubenoff R (2000). Skeletal muscle fiber quality in older men and women. Am J Physiol Cell Physiol 279, 611–618.

Galler S & Hilber K (1998). Tension/stiffness ratio of skinned rat skeletal muscle fibre types at various temperatures. Acta Physiol Scand 162, 119–126.

Galler S, Schmitt TL & Pette D (1994). Stretch activation, unloaded shortening velocity, and myosin heavy chain isoforms of rat skeletal muscle fibres. J Physiol 478, 513.

Geneva II, Cuzzo B, Fazili T & Javaid W (2019). Normal body temperature: A systematic review. Open Forum Infect Dis 6, 1–7.

Gilliver SF, Degens H, Rittweger J, Sargeant AJ & Jones DA (2009). Variation in the determinants of power of chemically skinned human muscle fibres. Exp Physiol 94, 1070– 1078.

Godt RE & Lindley BD (1982). Influence of temperature upon contractile activation and isometric force production in mechanically skinned muscle fibers of the frog. J Gen Physiol 80, 279– 297.

Goldman YE, McCray JA & Ranatunga KW (1987). Transient tension changes initiated by laser temperature jumps in rabbit psoas muscle fibres. J Physiol 392, 71–95.

Grosicki GJ, Gries KJ, Minchev K, Raue U, Chambers TL, Begue G, Finch H, Graham B, Trappe TA, Trappe S, Powers S & Hepple R (2021). Single muscle fibre contractile characteristics with lifelong endurance exercise. J Physiol 599, 14.

He ZH, Bottinelli R, Pellegrino MA, Ferenczi MA & Reggiani C (2000). ATP consumption and efficiency of human single muscle fibers with different myosin isoform composition. Biophys J 79, 945–961.

Hooft AM, Maki EJ, Cox KK & Baker JE (2007). An accelerated state of myosin-based actin motility. Biochemistry 46, 3513–3520.

Huxley AF (1957). Muscle structure and theories of contraction. Prog Biophys Biophys Chem 7, 255–318.

Huxley HE (1990). Sliding filaments and molecular motile systems. J Biol Chem 265, 8347– 8350.

Hvid LG, Ørtenblad N, Aagaard P, Kjaer M & Suetta C (2011). Effects of ageing on single muscle fibre contractile function following short-term immobilisation. J Physiol 589, 4745– 4757.

Jeon Y, Choi J, Kim HJ, Lee H, Lim JY & Choi SJ (2019). Sex- and fiber-type-related contractile properties in human single muscle fiber. J Exerc Rehabil 15, 537.

Kawai M (2003). What do we learn by studying the temperature effect on isometric tension and tension transients in mammalian striated muscle fibres? J Muscle Res Cell Motil 24, 127– 138.

Kawai M & Brandt PW (1980). Sinusoidal analysis: a high resolution method for correlating biochemical reactions with physiological processes in activated skeletal muscles of rabbit, frog and crayfish. J Muscle Res Cell Motil 1, 279–303.

Kawai M, Saeki Y & Zhao Y (1993). Crossbridge scheme and the kinetic constants of elementary steps deduced from chemically skinned papillary and trabecular muscles of the ferret. Circ Res 73, 35–50.

Kemp GJ, Meyerspeer M & Moser E (2007). Absolute quantification of phosphorus metabolite concentrations in human muscle in vivo by 31P MRS: a quantitative review. NMR Biomed 20, 555–565.

Kenny GP, Reardon FD, Zaleski W, Reardon ML, Haman F, Ducharme MB & Rear-Don ML (2003). Muscle temperature transients before, during, and after exercise measured using an intramuscular multisensor probe. J Appl Physiol 94, 2350–2357.

Knuth ST, Dave H, Peters JR & Fitts RH (2006). Low cell pH depresses peak power in rat skeletal muscle fibres at both 30°C and 15°C: Implications for muscle fatigue. J Physiol 575, 887–899.

Kojima H, Ishijima A & Yanagida T (1994). Direct measurement of stiffness of single actin filaments with and without tropomyosin by in vitro nanomanipulation. Proc Natl Acad Sci U S A 91, 12962–12966.

Krivickas LS, Dorer DJ, Ochala J & Frontera WR (2011). Relationship between force and size in human single muscle fibres. Exp Physiol 96, 539–547.

Kuhn HJ, Güth K, Drexler B, Berberich W & Rüegg JC (1979). Investigation of the temperature dependence of the cross bridge parameters for attachment, force generation and detachment as deduced from mechano-chemical studies in glycerinated single fibres from the dorsal longitudinal muscle of Lethocerus maximus. Biophys Struct Mech 6, 1–29.

Linari M, Bottinelli R, Pellegrino MA, Reconditi M, Reggiani C & Lombardi V (2004). The mechanism of the force response to stretch in human skinned muscle fibres with different myosin isoforms. J Physiol 554, 335–352.

Linari M, Caremani M, Piperio C, Brandt P & Lombardi V (2007). Stiffness and Fraction of Myosin Motors Responsible for Active Force in Permeabilized Muscle Fibers from Rabbit Psoas. Biophys J 92, 2476.

Lionikas A, Li M & Larsson L (2006). Human skeletal muscle myosin function at physiological and non-physiological temperatures. Acta Physiologica 186, 151–158.

Lovell SJ, Knight PJ & Harrington WF (1981). Fraction of myosin heads bound to thin filaments in rigor fibrils from insect flight and vertebrate muscles. Nature 293, 664–666.

Mebrahtu A, Smith IC, Liu S, Abusara Z, Leonard TR, Joumaa V & Herzog W (2024). Reconsidering assumptions in the analysis of muscle fibre cross-sectional area. J Exp Biol 227, 1–6.

Medler S (2019). Mixing it up: The biological significance of hybrid skeletal muscle fibers. J Exp Biol 22, 1–15.

Meijer JP, Jaspers RT, Rittweger J, Seynnes OR, Kamandulis S, Brazaitis M, Skurvydas A, Pišot R, Šimunič B, Narici M V. & Degens H (2015). Single muscle fibre contractile properties differ between body-builders, power athletes and control subjects. Exp Physiol 100, 1331– 1341.

Miller MS, Bedrin NG, Ades PA, Palmer BM & Toth MJ (2015). Molecular determinants of force production in human skeletal muscle fibers: Effects of myosin isoform expression and cross-sectional area. Am J Physiol Cell Physiol 308, 473–484.

Miller MS, Bedrin NG, Callahan DM, Previs MJ, Jennings ME, Ades PA, Maughan DW, Palmer BM, Toth MJ, II, Ades PA, Maughan DW, Palmer BM & Toth MJ (2013). Age-related slowing of myosin actin cross-bridge kinetics is sex specific and predicts decrements in whole skeletal muscle performance in humans. J Appl Physiol 115, 1004–1014.

Miller MS & Toth MJ (2013). Myofilament protein alterations promote physical disability in aging and disease. Exerc Sport Sci Rev 41, 93–99.

Miller MS, VanBuren P, LeWinter MM, Braddock JM, Ades PA, Maughan DW, Palmer BM & Toth MJ (2010). Chronic heart failure decreases cross-bridge kinetics in single skeletal muscle fibres from humans. J Physiol 588, 4039–4053.

Momb BA, Patino E, Akchurin OM & Miller MS (2022). Iron supplementation improves skeletal muscle contractile properties in mice with chronic kidney disease. Kidney 360 3, 843–858.

Momb BA, Patino E, Akchurin OM & Miller MS (2023a). Skeletal muscle single fiber force production declines early in juvenile male mice with chronic kidney disease. Physiol Rep 11, e15651.

Momb BA, Szabo GK, Mogus JP, Chipkin SR, Vandenberg LN & Miller MS (2023b). Skeletal muscle function is altered in male mice on low-dose androgen receptor antagonist or estrogen receptor agonist. Endocrinology 164, 1–11.

Mulieri LA, Barnes W, Leavitt BJ, Ittleman FP, LeWinter MM, Alpert NR & Maughan DW (2002). Alterations of myocardial dynamic stiffness implicating abnormal crossbridge function in human mitral regurgitation heart failure. Circ Res 90, 66–72.

Nelson CR, Debold EP & Fitts RH (2014). Phosphate and acidosis act synergistically to depress peak power in rat muscle fibers. Am J Physiol Cell Physiol 307, 939–950.

Ovejero JG, Fusi L, Park-Holohan SJ, Ghisleni A, Narayanan T, Irving M & Brunello E (2022). Cooling intact and demembranated trabeculae from rat heart releases myosin motors from their inhibited conformation. J Gen Physiol 154, e202113029.

Palmer BM, McConnell BK, Li GH, Seidman CE, Seidman JG, Irving TC, Alpert NR & Maughan DW (2004). Reduced cross-bridge dependent stiffness of skinned myocardium from mice lacking cardiac myosin binding protein-C. Mol Cell Biochem 263, 73–80.

Palmer BM, Suzuki T, Wang Y, Barnes WD, Miller MS & Maughan DW (2007). Two-state model of acto-myosin attachment-detachment predicts C-process of sinusoidal analysis. Biophys J 93, 760–769.

Palmer BM, Tanner BCW, Toth MJ & Miller MS (2013). An inverse power-law distribution of molecular bond lifetimes predicts fractional derivative viscoelasticity in biological tissue. Biophys J 104, 2540–2552.

Pathare N, Walter GA, Stevens JE, Yang Z, Okerke E, Gibbs JD, Esterhai JL, Scarborough MT, Gibbs CP, Sweeney HL & Vamdenborne K (2005). Changes in inorganic phosphate and force production in human skeletal muscle after cast immobilization. J Appl Physiol 98, 307–314.

Piazzesi G, Reconditi M, Koubassova N, Decostre V, Linari M, Lucii L & Lombardi V (2003). Temperature dependence of the force-generating process in single fibres from frog skeletal muscle. J Physiol 549, 93–106.

Rall JA & Woledge RC (1990). Influence of temperature on mechanics and energetics of muscle contraction. Am J Physiol 259, 197–203.

Ranatunga KW (2010). Force and power generating mechanism(s) in active muscle as revealed from temperature perturbation studies. J Physiol 588, 3657–3670.

Ranatunga KW (2018). Temperature effects on force and actin-myosin interaction in muscle: A look back on some experimental findings. Int J Mol Sci 19, 1–24.

Ranatunga KW, Sharpe B & Turnbull B (1987). Contractions of a human skeletal muscle at different temperatures. J Physiol 390, 383–395.

Reiser PJ, Welch KC, Suarez RK & Altshuler DL (2013). Very low force-generating ability and unusually high temperature dependency in hummingbird flight muscle fibers. J Exp Biol 216, 2247–2256.

Schneider CA, Rasband WS & Eliceiri KW (2012). NIH Image to ImageJ: 25 years of image analysis. Nat Methods 9, 671–675.

Seitz LB & Haff GG (2016). Factors modulating post-activation potentiation of jump, sprint, throw, and upper-body ballistic performances: A systematic review with meta-analysis. Sports Medicine 46, 231–240.

Smith IC & Herzog W (2023). Assumptions about the cross-sectional shape of skinned muscle fibres can distort the relationship between muscle force and cross-sectional area. J Appl Physiol 135, 1036–1040.

Stephenson G (2001). Hybrid skeletal muscle fibres: A rare or common phenomenon? Clin Exp Pharmacol Physiol 28, 692–702.

Stienen GJM, Kiers JL, Bottinelli R & Reggiani C (1996). Myofibrillar ATPase activity in skinned human skeletal muscle fibres: fibre type and temperature dependence. J Physiol 493, 299– 307.

Stienen GJM, Rooselmalen MCM, Wilson MGA & Elzinga G (1990). Depression of force by phosphate in skinned skeletal muscle fibers of the frog. Am J Physiol 259, 359–47.

Sundberg CW, Hunter SK, Trappe SW, Smith CS & Fitts RH (2018). Effects of elevated H+ and Pi on the contractile mechanics of skeletal muscle fibres from young and old men: implications for muscle fatigue in humans. J Physiol 596, 3993–4015.

Tanner BCW, Daniel TL & Regnier M (2007). Sarcomere lattice geometry influences cooperative myosin binding in muscle. PLoS Comput Biol 3, 1195–1211.

Toth MJ, Miller MS, Callahan DM, Sweeny AP, Nunez I, Grunberg SM, Der-Torossian H, Couch ME & Dittus K (2013). Molecular mechanisms underlying skeletal muscle weakness in human cancer: Reduced myosin-actin cross-bridge formation and kinetics. J Appl Physiol 114, 858–868.

Troiano RP, Berrigan D, Dodd KW, Mâsse LC, Tilert T & Mcdowell M (2008). Physical activity in the United States measured by accelerometer. Med Sci Sports Exerc 40, 181–188.

Tyska MJ & Warshaw DM (2002). The myosin power stroke. Cell Motil Cytoskeleton 51, 1–15.

Wakabayashi K, Sugimoto Y, Tanaka H, Ueno Y, Takezawa Y & Amemiya Y (1994). X-ray diffraction evidence for the extensibility of actin and myosin filaments during muscle contraction. Biophys J 67, 2422–2435.

Walcott S, Warshaw DM & Debold EP (2012). Mechanical coupling between myosin molecules causes differences between ensemble and single-molecule measurements. Biophys J 103, 501–510.

Wang G & Kawai M (1997). Force generation and phosphate release steps in skinned rabbit soleus slow-twitch muscle fibers. Biophys J 73, 878.

Wang G & Kawai M (2001). Effect of temperature on elementary steps of the cross-bridge cycle in rabbit soleus slow-twitch muscle fibres. J Physiol 531, 219–234.

Wilson CJ, Nunes JP & Blazevich AJ (2025). The effect of muscle warm-up on voluntary and evoked force-time parameters: A systematic review and meta-analysis with meta-regression. J Sport Health Sci 14, 101024.

Zhao Y & Kawai M (1993). The effect of the lattice spacing change on cross-bridge kinetics in chemically skinned rabbit psoas muscle fibers. II. Elementary steps affected by the spacing change. Biophys J 64, 197–210.

Zhao Y & Kawai M (1994). Kinetic and thermodynamic studies of the cross-bridge cycle in rabbit psoas muscle fibers. Biophys J 67, 1655–1668.

